# Highly multiplexed imaging recovers immune and metabolic niches in multiple myeloma associated with disease progression and bone involvement

**DOI:** 10.1101/2025.09.08.674473

**Authors:** Ingrid Aass Roseth, Lukas Hatscher, Chiara Schiller, Håkon Skjalg Selland Johnstuen, Tobias S. Slørdahl, Håkon Hov, Denis Schapiro, Therese Standal

## Abstract

Multiple myeloma (MM) is a cancer arising from genetically aberrant plasma cells (PCs) that remain dependent on the bone marrow (BM) microenvironment for disease establishment and progression. Here, we spatially mapped BM biopsies from patients with MM, smoldering MM and monoclonal gammopathy of undetermined significance by imaging mass cytometry. We found that PCs near bone surfaces were quiescent and demonstrated a distance–to-bone-dependent upregulation of IL-32 and HIF1α in patients with bone disease, consistent with their role in promoting bone loss in MM. We further identified two distinct PC neighborhoods termed PC_OXPHOS characterized by focal PC growth, enrichment of endothelial cells and elevated oxidative phosphorylation; and PC_MYELOID, characterized by PCs interspersed with immune cells and featuring a glycolytic phenotype. Spatial interactions between PCs and immune correlated with shorter progression free survival. Notably, a strong neighbor preference between PCs and CD4^+^ T cells was an independent predictor for disease progression. Our work underscores spatial context as a key factor in understanding MM pathogenesis and the potential for spatial analyses to improve MM risk stratification.

**Significance:** This study demonstrates that MM cells in different neighborhoods experience unique metabolic conditions and immune environments, and that neighbor preference of CD4^+^ T cells and plasma cell is associated with disease progression. These features were not captured by non-spatial metrics, highlighting the value of spatially resolved analyses in uncovering pathophysiological mechanisms of MM.

## Introduction

Multiple myeloma (MM) is the second most common hematologic cancer and is caused by the accumulation of malignant plasma cells (PCs) in the bone marrow.^1^ MM is preceded by the premalignant conditions monoclonal gammopathy of unknown significance (MGUS) and smoldering myeloma (SMM). The premalignant PCs have genetic aberrations in common with the tumor PCs, and progression to MM happens when the PCs acquire secondary genetic mutations.^1^ Development of MM is also accompanied by changes in the TME leading to immune suppression and, for most patients, a severe osteolytic bone disease.^2,3^

MM is a heterogenous disease, and while some patients have a poor prognosis with survival time of less than two years after diagnosis, others survive for more than a decade.^4^ The risk for rapid disease progression can, to some extent, be predicted based on the genetic makeup of the PC and by serum biomarkers.^5^ The old staging system has been updated in recent years.^4,6^ However, the system is still not perfect and does not take the contribution of the TME on disease progression into account.

BM aspirates are routinely obtained from MM patients at diagnosis, and high-throughput techniques such as multiplex flow cytometry and single-cell RNA sequencing (scRNA-seq) have significantly advanced our understanding of the TME in MM.^7–9^ In terms of prognostication, it was recently shown that that the immune composition in the BM can predict overall survival in MM patients irrespective of cytogenetics risk profile.^10^ However, important cell types, such as bone cells, stromal cells and adipocytes are missing in BM aspirates, and the data provides no information about cell-cell interactions or spatial aspects of tumor growth or immune cell infiltration. Addressing these shortcomings, a few recent studies have examined BM biopsies from MM either using cyclic immunofluorescence or multiplexed immunohistochemistry.^11,12^ However, a restricted numbers of protein markers in the previous studies give only a limited understanding of BM tissue structures and cellular interactions.

To achieve high-resolution spatial profiling, we employed imaging mass cytometry (IMC) with a 35-antibody panel on trephine BM biopsies from patients with newly diagnosed untreated MM, MGUS and SMM. We identified fourteen different cell types and assessed their organization into neighborhoods based on cell type-specific, functional and metabolic features. We identified a distance to bone-dependent PC signature that is associated with bone disease (BD), and that the BM tissue is organized into neighborhoods with distinct metabolic profiles. Focal growth of PCs distinguishes MM from SMM and MGUS, and PCs in focal micro clusters have a distinct metabolic profile with active mitochondrial metabolism and increased amino acid uptake. We also demonstrate for the first time that a spatial feature, CD4^+^ T cells and PC neighbor interactions, can provide prognostic information in MM.

## Results

### High-dimensional imaging defines the cellular composition of MM

To spatially characterize the MM TME, we stained biopsies from newly diagnosed, untreated MM, MGUS and SMM patients (Supplementary Table 1) with a panel of antibodies which enabled identification of the main cell types in the BM and their cellular metabolism^13^, proliferation and activation (Fig. 1A, Supplementary Table 2, Supplementary Fig.1, Supplementary Fig. 2A). To ensure high confidence in cell type assignment and minimize misclassification, we employed a stringent classification approach (Supplement Fig. 2B). Cell identity was verified by inspection of marker gene expression (Supplementary Fig. 3) and by overlaying signal intensities from raw IMC images with phenotype labels (Figure 1B). This approach enabled phenotyping of 1,007,127 cells into 14 distinct cell types (Fig. 1C, Supplementary Fig. 4A). As the antibody panel was tailored towards functional markers, a proportion of cells remained unclassified as expected (Fig. 1C).

**Fig. 1.**
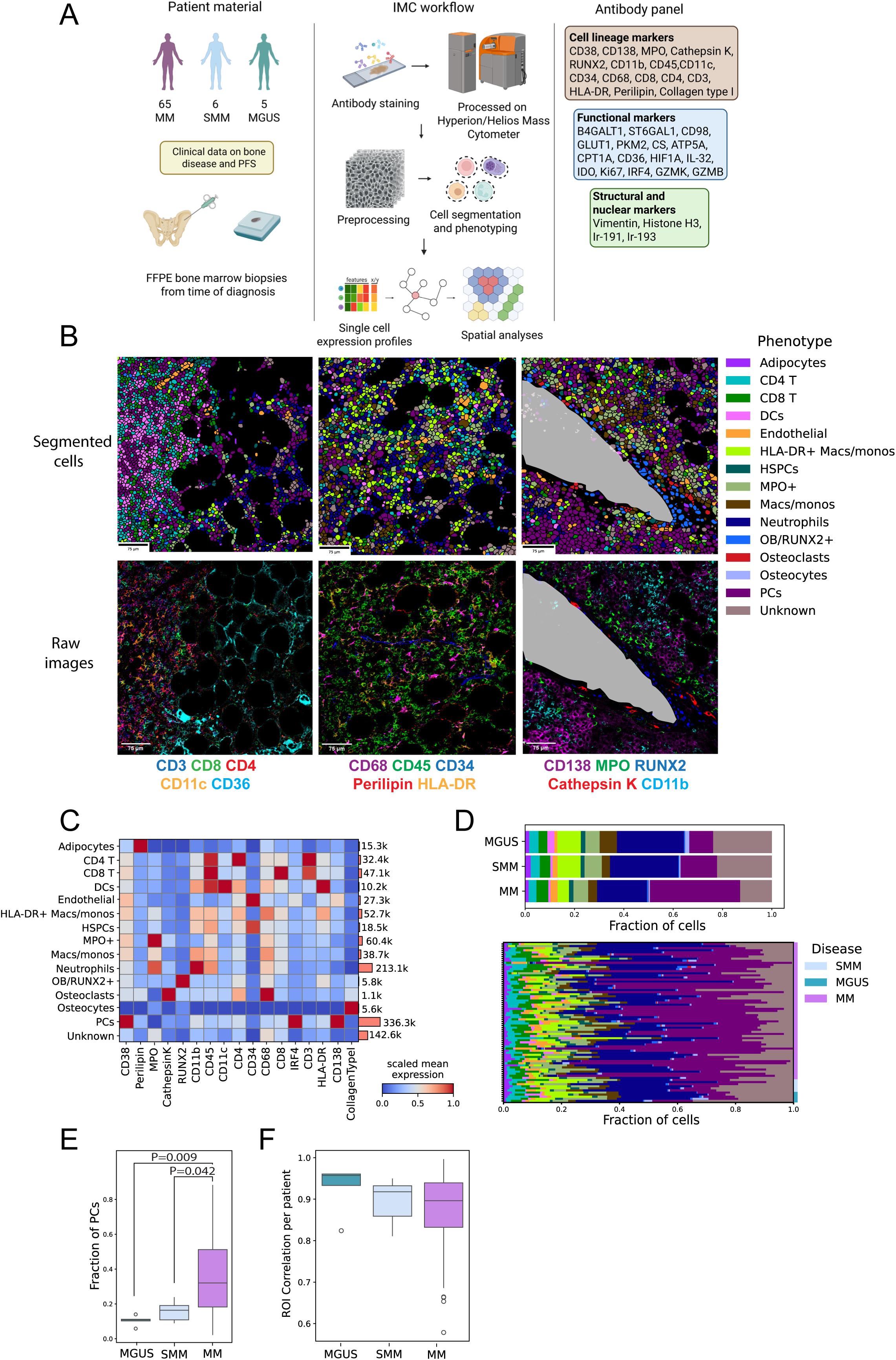
High-dimensional imaging defines the cellular composition of MM. **(A)** Overview of the patient cohort, imaging and data analysis workflow. Illustration created using Biorender.com **(B)** Phenotype plots with segmentation mask boundaries (top) with corresponding raw IMC images (bottom) of selected markers. Bone surfaces were manually traced and illustrated in grey. **(C)** Heatmap showing scaled mean expression of cell type and cell lineage specific markers across all cell types. **(D)** Cumulated cell type composition for MM (N= 65), SMM (N=6) and MGUS (N=5) patients (top plot) and cell type composition for each individual patient (bottom plot). **(E)** Frequencies of plasma cells (PCs) per image. The median is indicated by a horizontal solid line, whiskers extending to the most extreme data points within 1.5 times the interquartile range from the quartiles. Points beyond the whiskers represent outliers. P-values were determined by Kruskal-Wallis with Dunn’s post-hoc test with Bonferroni correction. Non-significant comparisons are omitted from the plot. **(F)** Spearman correlation coefficients of cell type composition of two ROIs of individual patients. The median is indicated by a horizontal solid line, whiskers extending to the most extreme data points within 1.5 times the interquartile range from the quartiles. Points beyond the whiskers represent outliers.

The relative abundance of cell types in MGUS, SMM and MM reflected disease biology with an increased abundance of PCs from MGUS and SMM to MM (Fig. 1D and E, Supplementary Fig. 4B). Although average correlations in cellular composition across disease groups were high (Fig. 1F), regions of interest (ROIs) from the same biopsy showed greater variability in MM than in SMM or MGUS, reflecting increased heterogeneity in diseased BM. The significant correlation between the percentage of PCs estimated by the pathologist in the biopsy and the percentage derived from our analyses on the same biopsy (Supplementary Fig. 4C), suggests that the ROIs were representative for most patients. However, since the two ROIs per patient represent only a small fraction of the total biopsy, some outliers were present (Supplementary Fig. 4D).

### Quiescent PCs are located near bone surfaces

The endosteal niche is a specialized microenvironment within the bone marrow, located near the inner surface of bones (endosteum), where hematopoietic stem cells (HSCs) reside.^14^ It plays a crucial role in maintaining stem cell quiescence, self-renewal, differentiation, and has been shown to contribute to the survival of quiescent stem-like MM cells in mice.^15–17^ We calculated the individual cells’ distance to the nearest bone (Fig. 2A) and found that overall, the different immune cell types showed similar distances to bone, while as expected osteoclasts (OCs), osteocytes and osteoblasts (OBs), which are denoted in our study as OB/RUNX2^+^ cells, were found at or near bone surfaces (Fig. 2B). Strikingly, when we examined if PCs location relative to bone impacted the cells’ phenotype, we found that the fraction of non-proliferating PCs was enriched towards bone surfaces (Fig. 2C). The trend of quiescent PCs for this niche was similar for cells in all disease groups, supporting the notion that this is a general feature of human PCs rather than a malignancy-associated phenomenon (Fig. 2C).

**Fig. 2.**
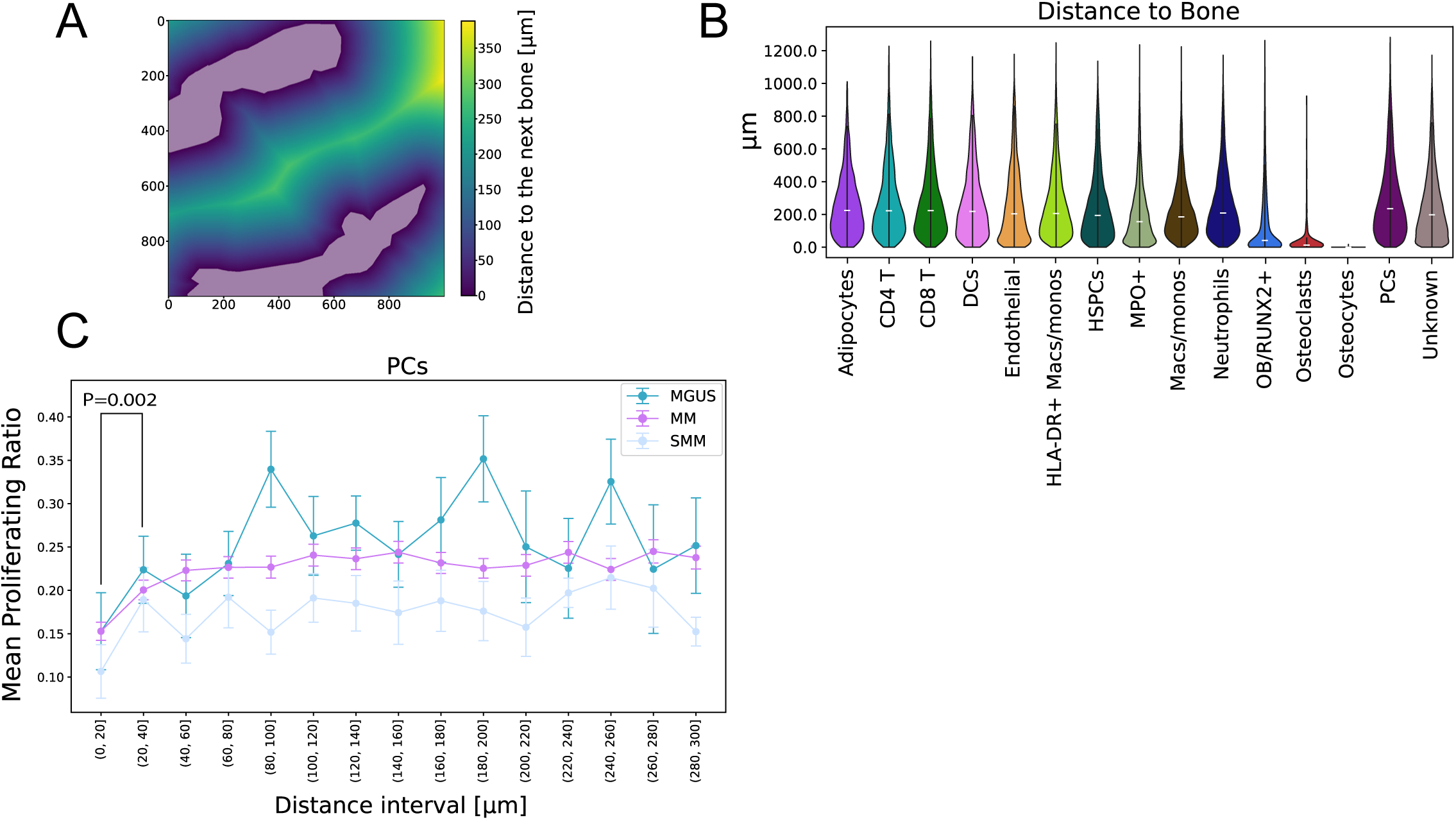
Quiescent PCs are located near bone surfaces. **(A)** Illustration showing pixel-wise calculation of the distance [µm] to the next bone mask (grey). **(B)** Distance to nearest bone [µm] for all cell types. The median is indicated by a horizontal solid line. **(C)** Proliferation ratio by distance to bone surface [µm], calculated as Ki67^+^ PCs divided by all PCs in 20 µm increments. P-value was determined by Mann-Whitney-U-test. Non-significant comparisons are omitted from the plot.

### Spatial profiling of PCs’ marker expression reveals a bone-proximal expression signature linked to bone disease

Nearly all MM patients develop osteolytic lesions at diagnosis or during disease progression, driven by a complex interplay between malignant cells and the tumor microenvironment (TME) that increases osteoclast (OC) number and activity while reducing and impairing osteoblasts (OBs).^18,19^ To study if the location of PCs relative to bone and bone cells is associated with bone disease (BD), we grouped MM patients (N=65) into patients with BD (MM_BD) (N=55) and patients without BD (MM_noBD) (N=10) upon time of diagnosis (Fig. 3A, Supplementary Table 3).

**Fig. 3.**
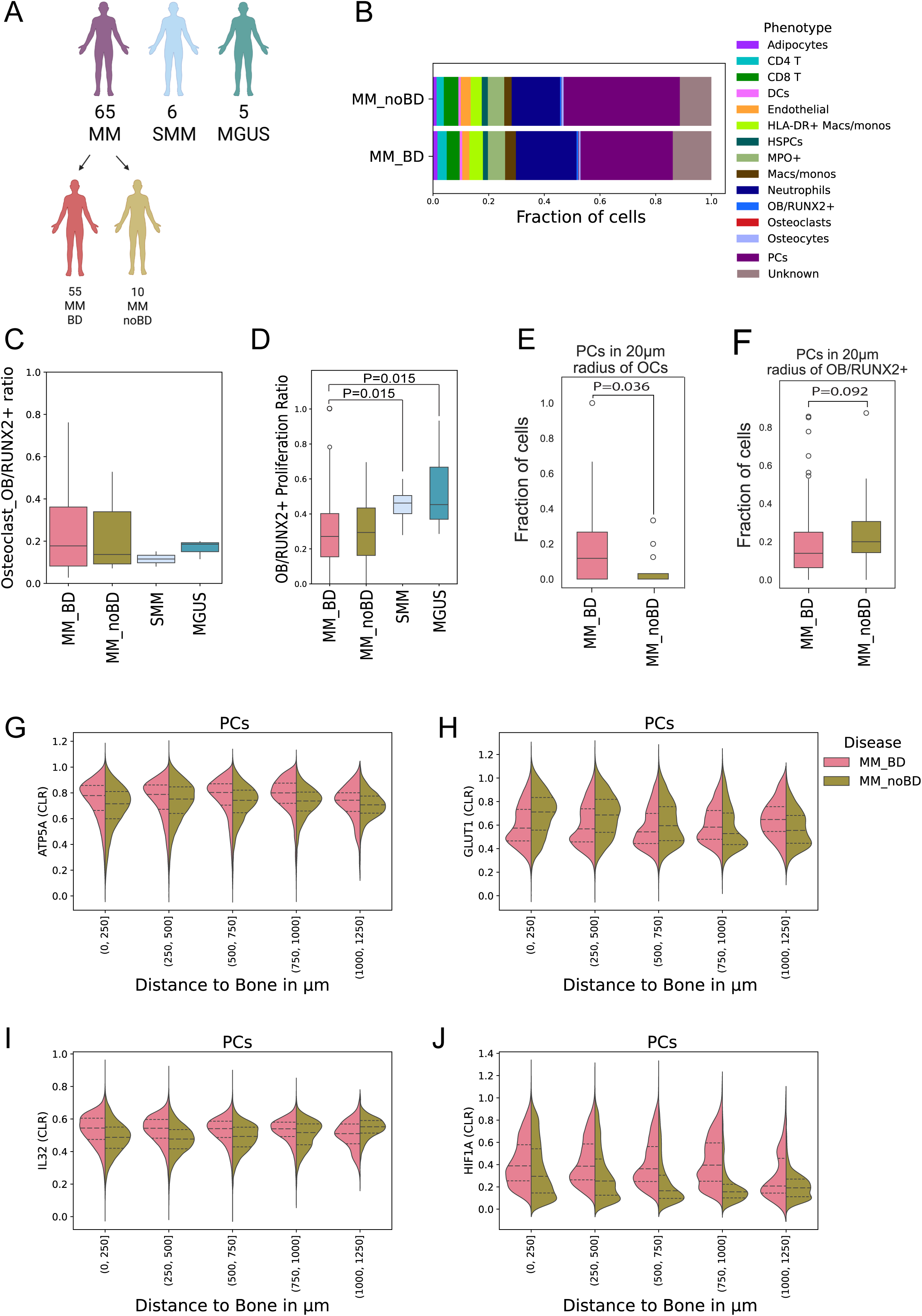
Spatial profiling of PCs’ marker expression reveals a bone-proximal expression signature linked to bone disease. **(A)** Myeloma patients (MM, N= 65) were grouped according to presence or absence of bone disease as evaluated by whole-body low dose CT. Patients with bone disease: MM_BD, N=55, patients without bone disease: MM_noBD, N=10, SMM (N=6) and MGUS (N=5). Illustration created using Biorender.com **(B)** Cell type abundances presented as fractions of total number of cells in the cohorts. **(C)** Ratio of osteoclasts to OB/RUNX2^+^ cells per image. The median is indicated by a horizontal solid line, whiskers extending to the most extreme data points within 1.5 times the interquartile range from the quartiles. Points beyond the whiskers represent outliers. The differences between groups were not significant (Kruskal-Wallis test). **(D)** Proliferation ratio of osteoblasts (OB)/RUNX2+ cells, calculated as the number of proliferating (Ki67^+^) OB/RUNX2^+^ cells to all OB/RUNX2^+^ cells. The median is indicated by a horizontal solid line, whiskers extending to the most extreme data points within 1.5 times the interquartile range from the quartiles. Points beyond the whiskers represent outliers. P-values were determined by Kruskal-Wallis with Dunn’s post-hoc test with Bonferroni correction. Non-significant comparisons are omitted from the plot. **(E)** Proportion of PCs within a 20 µm radius of osteoclasts (OCs) as median per image. The median in each cohort is indicated by a horizontal solid line, whiskers extending to the most extreme data points within 1.5 times the interquartile range from the quartiles. Outliers are shown as individual points. P-value was determined by Mann-Whitney-U-Test. **(F)** Proportion of PCs within a 20 µm radius of OB/RUNX2^+^ cells as median per image. The median in each cohort is indicated by a horizontal solid line, whiskers extending to the most extreme data points within 1.5 times the interquartile range from the quartiles. Outliers are shown as individual points. P-values determined by Mann-Whitney-U-Test. CLR-normalized mean expression of ATP5A **(G)**, GLUT1 **(H)**, IL-32 **(I)** and HIF1a **(J)** in PCs, stratified into five bins based on distance to bone. Dashed lines in the violin plots represent 0.25, 0.5 and 0.75 quartiles.

Cell type abundances were overall similar in both groups (Fig. 3B, Supplementary Fig. 5A) and there were no significant differences in numbers of OCs per OB/RUNX2^+^ cells between the groups (Fig. 3C). In contrast, MM patients had reduced frequencies of proliferating OB/RUNX2^+^ compared with SMM and MGUS (Fig. 3D), in support of the known osteoblast suppression in myeloma^20^, while there was no difference in OB/RUNX2^+^ cell proliferation or their frequency in patients with or without BD (Fig. 3D, Supplementary Fig. 5A).

Since non-spatial metrics did not distinguish patients with and without bone disease (BD), we quantified the spatial abundance of plasma cells (PCs) near bone cells. A high abundance of PCs was observed within the immediate vicinity (20 µm) of OCs in patients with BD, a phenomenon that was nearly absent in patients without BD (Fig. 3E), and independent of the overall PC abundance (Fig. 3B, Supplementary Fig. 5A). These findings support the idea that direct interactions between MM cells and OCs contribute to bone destruction, consistent with previous reports.^21^ In contrast, PC abundance near osteoblasts (OBs)/RUNX2⁺ cells tended to be lower in patients with BD, although this difference did not reach statistical significance (Fig. 3F).

We next investigated whether proximity to OCs influences expression of specific functional markers related to metabolism, immune response or proliferation in PCs. Given the scarcity of OCs in the biopsies, we used distance to bone as a proxy measure.

In patients with BD, PCs exhibited a general tendency of elevated ATP5A expression, a key marker of OXPHOS, regardless of their proximity to bone surfaces (Fig. 3G). In contrast, GLUT1 expression, indicating active glycolysis, displayed a distance dependent gradient and the difference between cohorts diminished with increasing distance (Fig. 3H). There was no difference among the groups using HistoneH3 as a control (Supplementary Fig. 5B). Of note, analysis revealed that the expression of IL-32, a cytokine implicated in bone disease^22^, showed a trend of increased expression in BD compared with noBD in areas close to the bone, which progressively diminished with increasing distance (Fig. 3I). A similar trend was observed for heat-inducible factor 1 alpha (HIF1a), which has been shown to promote IL-32 expression in malignant PCs^22^ (Fig. 3J). In summary, functional properties of PCs such as metabolic activity and cytokine expression varied depending on their location relative to bone surfaces. Our results support that crosstalk between bone cells and metabolically and functionally altered PCs could impact the development of bone disease in MM or vice versa.

### The BM is organized into spatially resolved neighborhoods defined by cellular composition and cellular activation states

A cellular neighborhood refers to a group of cells that frequently occur together in a tissue, suggesting a shared underlying function.^23^ To identify BM neighborhoods (NBHs) driven by both cell type composition and functional marker expression that collectively shape their spatial organization, we employed unsupervised neighborhood analysis with CellCharter.^24^ While some methods solely rely on cell type labels and result in identification of cellular neighborhoods^25^, CellCharter also takes functional markers into account, making it particularly suitable for our antibody panel design (Fig. 4A). Following a cluster stability analysis and manual review of top hits, we selected the number of NBHs to be 9 (Supplementary Fig. 6A).

**Fig. 4.**
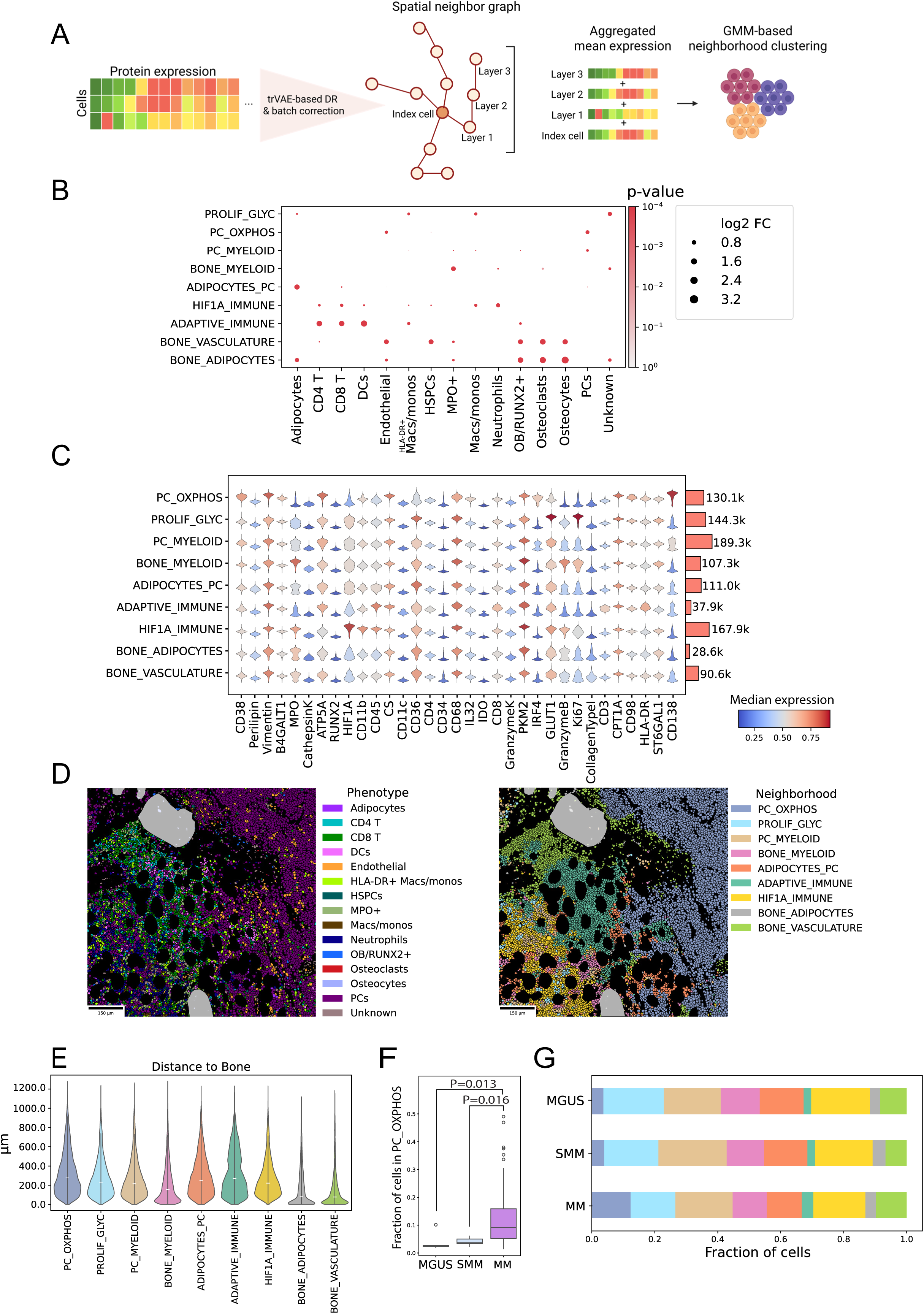
The BM is organized into spatially resolved neighborhoods defined by cellular composition and cellular activation states. **(A)** Schematic overview of CellCharter’s workflow to generate neighborhood labels: Dimensionality reduction to 10 features and batch correction was performed using CellCharter’s adaptation of scarches’ trVAE. After creating a spatial neighbor graph using squidpy, CellCharter aggregated features of 3 layers of cells. Neighborhood labels were then inferred using GMM-based spatial clustering. Illustration created using Biorender.com **(B)** Phenotype enrichment plot showing the relative enrichment of cell types (log2 Fold Change) among neighborhoods. 250-fold permutation of neighborhood labels is used to infer empirical P-values. **(C)** Arcsinh-transformed median protein marker expression of cells in different neighborhoods**. (D)** Phenotype-(left) and neighborhood (right) plots with segmentation boundaries displaying representative phenotype- and neighborhood mappings. Manually traced bone surfaces illustrated in grey. **(E)** Distance to bone [µm] for all cells in different neighborhoods. Median is illustrated in the violin plots by a horizontal line. **(F)** Fraction of cells in each image located in PC_OXPHOS neighborhood. The median in each group is indicated by a horizontal solid line, whiskers extending to the most extreme data points within 1.5 times the interquartile range from the quartiles. Outliers are shown as individual points. P-values determined by Kruskal-Wallis with Dunn’s post-hoc test with Bonferroni correction. Non-significant comparisons are omitted from the plot. **(G)** Cumulated neighborhood composition for MM, SMM and MGUS.

Phenotype enrichment analysis revealed that PCs were predominantly present in two NBHs: One termed PC_OXPHOS, due to the enrichment of PCs and markers associated with OXPHOS and the second PC_MYELOID, which showed enrichment of both PCs and myeloid cells (Fig. 4B-D). PCs were also to some extent present in the ADIPOCYTES_PC NBH, that apart from PCs were enriched in adipocytes and CD8^+^ T cells (Fig. 4B and D).

Two immune cell-enriched NBHs termed ADAPTIVE_IMMUNE, and HIF1A_IMMUNE were identified. The ADAPTIVE_IMMUNE NBH was dominated by CD4^+^ T cells, CD8^+^ T cells, DCs and HLA-DR^+^ macrophages/monocytes (Fig. 4B and D). High expression of granzyme K (GZMK) and human leukocyte antigen-DR (HLA-DR), suggests that this NBH consisted of activated T cells and antigen presenting cells (Fig. 4C). Of note, the ADAPTIVE_IMMUNE NBH highly expressed indoleamine 2,3 dioxygenase (IDO), reflecting immune regulation typically observed post-activation or during chronic active immune responses^26^ (Fig. 4C). HIF1A_IMMUNE was dominated by myeloid cells such as neutrophils and HLA-DR^−^ macrophages/monocytes (Fig. 4B and D). High expression of HIF1a suggests hypoxic conditions in this niche (Fig. 4C). A third NBH, termed PROLIF_GLYC was predominantly characterized by GLUT1 and Ki67 expression and enriched for monocytes/macrophages (Fig. 4B-D).

Three NBHs termed BONE_VASCULATURE, BONE_MYELOID and BONE_ADIPOCYTE were enriched for bone cells (OB/RUNX2^+^ cells, OCYs and OCs) with varying co-enrichment of other cell types (Fig. 4B and D). Among these, BONE_MYELOID was located further from bone surfaces than BONE_ADIPOCYTES and BONE_VASCULATURE (Fig. 4E), suggesting that the latter two NBHs may constitute the ‘endosteal niche’.^14^

Overall, most NBHs were of similar size, varying between 90 000 to 190 000 cells, while BONE_ADIPOCYTES and ADAPTIVE_IMMUNE were smaller, encompassing about 28 000 and 40 000 cells, respectively (Fig. 4C). There was an increased abundance of PC_OXPHOS in the MM group compared with MGUS and SMM (Fig. 4F and G), supporting a change in growth pattern of PCs upon malignancy. Similar to what we observed for cell type composition, there was substantial inter-individual variation in neighborhood abundances (Supplementary Fig. 6B).

### Distinct metabolic profiles of PCs depend on their localization in different neighborhoods

MM is characterized by a focal growth pattern of tumor cells at the macroscopic level in the skeleton.^27,28^ At the microscale, tumor cells in the BM also exhibit a patchy distribution, in contrast to the more diffuse PC infiltration typically seen in precursor states.^11,12^ Notably, the two PC-enriched neighborhoods, PC_OXPHOS and PC_MYELOID, displayed distinct spatial growth patterns. PC_OXPHOS formed dense, focal aggregates, while PC_MYELOID was more dispersed and interspersed with immune cells, resembling the growth patterns more commonly observed in pre-malignant conditions (Fig. 5A). To quantify these differences, we assessed the spatial cohesion of all neighborhoods by evaluating the stability of their connected components as we progressively increased the minimum cell threshold required to define a connected structure within the spatial graph (Fig. 5B). PC_OXPHOS remained connected far longer than other neighborhoods, with nearly 60% of its cells still forming connected components at a threshold of 200 cells (Fig. 5C), indicating strong aggregation. In contrast, PC_MYELOID was much less stable, with fewer than 15% of its cells connected at the same threshold. This aggregation phenotype of PC_OXPHOS was predominantly observed in MM patients, suggesting an association with malignancy (Fig. 5D).

**Fig 5.**
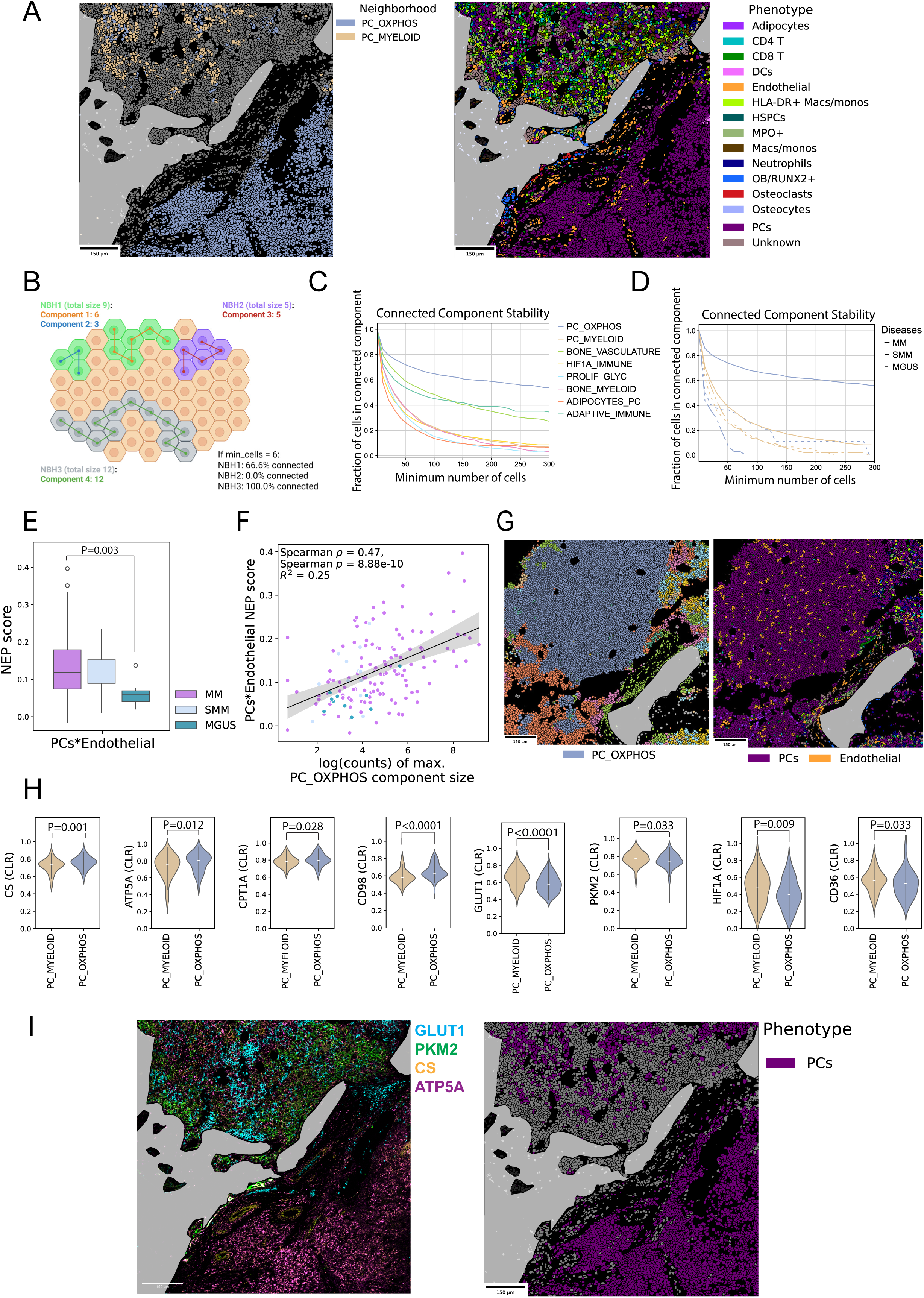
Distinct metabolic profiles of PCs depend on their localization in different neighborhoods. **(A)** Neighborhood- (left) and phenotype (right) plots with segmentation boundaries. Manually traced bone surfaces illustrated in grey. **(B)** Illustration of connected components. A connected component is defined as a specific number of cells from the same neighborhood directly connected on the spatial neighbor graph given a specific threshold. The number of these connected cells is then related to the total number of cells from that neighborhood. Connected component stability was assessed by calculating the fraction of cells residing inside a connected component by increasing the minimal number of cells required to form a connected component. Illustration created using Biorender.com **(C+D)** Connected component stability across all neighborhoods showing the percentage (y-axis) of cells in each neighborhood still connected at various sizes of connected components (x-axis) in the whole cohort (left) and when divided into subgroups of MM, SMM and MGUS patients (right). **(E)** Pair-wise neighbor preference (NEP) scores between PCs and endothelial cells (ECs). NEP scores are normalized per image. The median in each cohort is indicated by a horizontal solid line, whiskers extending to the most extreme data points within 1.5 times the interquartile range from the quartiles. Outliers are shown as individual points. P-values determined by Kruskal-Wallis with Dunn’s post-hoc test with Bonferroni correction. Non-significant comparisons are omitted from the plot. **(F)** Spearman correlation coefficients of the normalized pair-wise NEP score of PCs*ECs and the max component size of PC_OXPHOS per image. Spearman’s ρ = 0.47, p = 8.88*10^−10^ from correlation analysis, R² = 0.25 calculated from linear regression. **(G)** Neighborhood- (left) and phenotype (right) plots with segmentation boundaries to illustrate vascularization of PC_OXPHOS neighborhood. Manually traced bone surfaces are illustrated in grey. **(H)** CLR-normalized mean expression of metabolism-related markers per image of PCs from connected components (60 cells) of PC_MYELOID and PC_OXPHOS. Median is illustrated in the violinplots by a horizontal line. P-values determined by Mann-Whitney-U-Test. **(I)** Raw IMC image highlighting the expression of GLUT1, PKM2, CS and ATP5A (left) and an associated phenotype spatial plot showing locations of PCs (right).

The phenotype enrichment analysis revealed that PC_OXPHOS contained more Endothelial cells (ECs) than other NBHs (Fig. 4B). To study interactions between PCs and ECs in more detail, we performed permutation based pairwise neighbor preference analysis using COZI.^29^ COZI evaluates whether two cells have a spatial preference for each other compared to a permuted spatial randomness. Higher neighbor preference of PCs to ECs seemed to be associated with progression from MGUS to SMM and MM (Fig. 5E). Moreover, the EC-PC preference score positively correlated with the maximum component size of PC_OXPHOS (Fig. 5F and G). These findings indicate that PC_OXPHOS aggregates are highly vascularized, further supporting the notion that this NBH exhibits a respiratory metabolic phenotype.

To evaluate cellular metabolism, our antibody panel included markers important for regulating the main metabolic pathways (Supplementary Fig. 2A).^13^ To further examine if the OXPHOS phenotype in PC_OXPHOS was related to the focal growth pattern, we evaluated marker expression in PCs within a connected component of at least 60 cells to ensure analysis is performed on aggregated cells (Fig. 5H). Indeed, PCs from the PC_OXPHOS NBH showed signs of active mitochondrial metabolism with a higher expression of CS, ATP5A, CPT1A and increased amino acid transport through increased expression of CD98 compared with PCs from PC_MYELOID (Fig. 5H and I). In contrast, the expression of HIF1a, GLUT1, PKM2 and CD36 were higher in PCs from PC_MYELOID (Fig. 5H and 4I), which indicates a less vascularized area with a preference for anaerobic glycolysis. Taken together, these results support that PCs in the two identified cancer cell-dominated NBHs have distinct metabolic profiles: PCs in focal micro clusters prefer OXPHOS/FAO supported by increased vascularization while PCs with interspersed growth pattern in the PC_MYELOID rely more on glycolysis-dependent metabolism.

### Focal growth of PCs is associated with immune cell avoidance

To decipher general interaction patterns between PC and immune cells in detail, we grouped all immune cells into one category (CD4^+^ T, CD8^+^ T, DCs, macrophages/monocytes and neutrophils) and applied COZI^29^ (Fig. 6A). In premalignant samples, we found that all samples shared a weak-to moderate neighbor preference of immune cells to PCs, while MM patients displayed heterogeneous patterns, i.e. either stronger preference or avoidance, implying immune cell-PC interaction or exclusion, respectively (Fig. 6A-C). This was also evident when the neighbor preference scores between immune cell subtypes and PCs were calculated individually (Supplementary Fig. 7). This characteristic seemed to be associated with the spatial growth pattern of PCs as the immune cell-PC neighbor preference negatively correlated with the maximum component size of PC_OXPHOS (rho = -0.67) (Fig. 6D). This suggests that PCs in large PC_OXPHOS aggregates might be protected from immune cells.

**Fig. 6.**
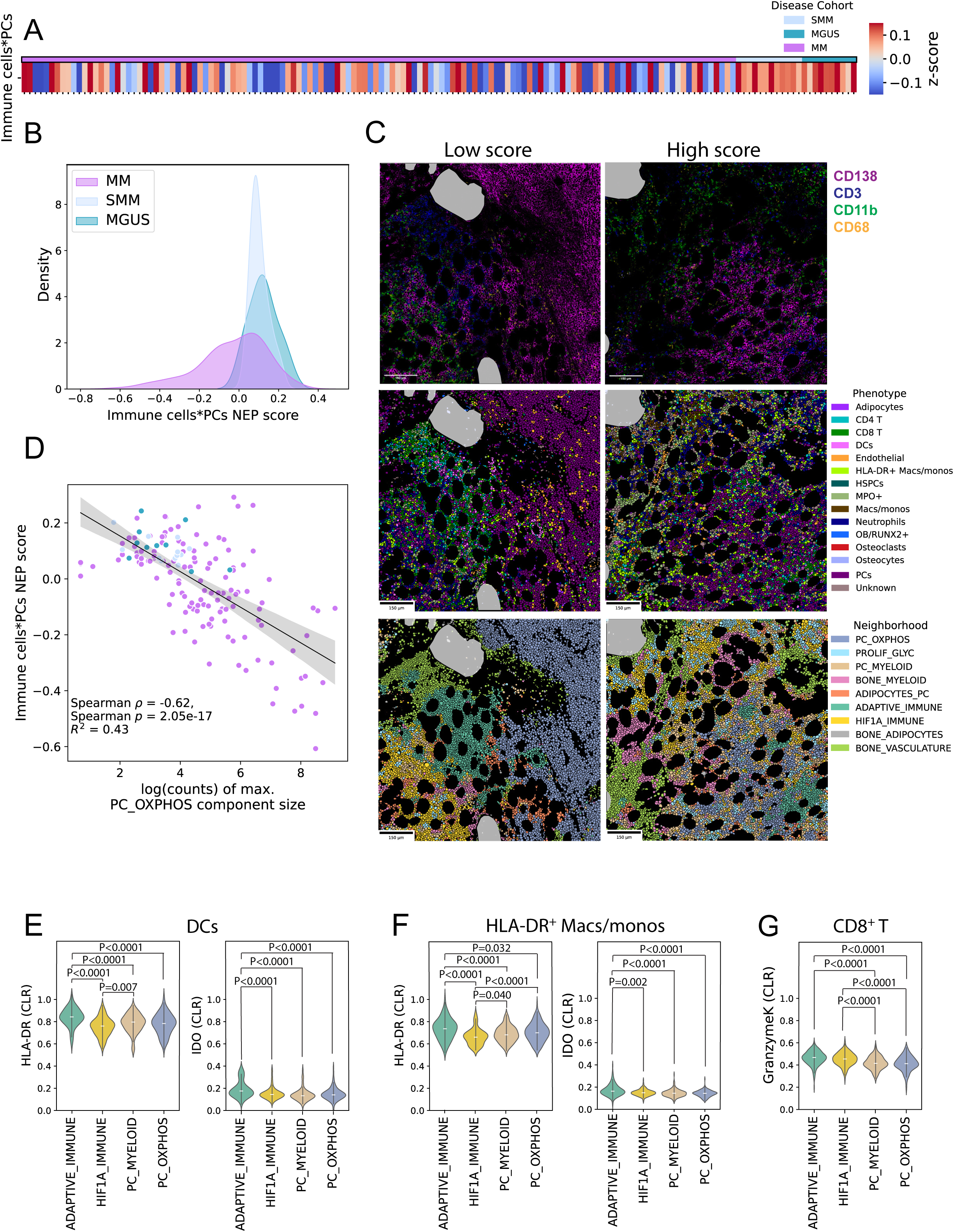
Focal growth of PCs is associated with immune cell avoidance and neighborhood context influences immune cell activation marker profile in MM. **(A)** Neighbor preference (NEP) scores between PCs and immune cells obtained by grouping CD4 T, CD8 T, DCs, HLA-DR^+^/^−^ Macs/monos and neutrophils. NEP scores are normalized per image. **(B)** Density plot of normalized pairwise NEP scores from (A) **(C)** Raw IMC images (top) highlighting ROIs with low and high normalized NEP scores linked to the corresponding spatial phenotype (middle) and neighborhood (bottom) plots. Manually traced bone surfaces are illustrated in grey. **(D)** Spearman correlation coefficients of normalized pair-wise NEP scores from (A) and the max component size of PC_OXPHOS per image. Spearman’s ρ = -0.62, p = 2.05*10^−17^, R² = 0.43 calculated from linear regression. **(E-F)** CLR-normalized mean expression per image of HLA-DR and IDO in **(E)** DCs and **(F)** HLA-DR^+^ Macs/monos in immune- and cancer-related neighborhoods. **(G)** CLR-normalized mean expression per image of GZMK in CD8^+^ T in immune- and cancer-related neighborhoods. Median is illustrated in the violinplots by a horizontal line. P-values determined by Kruskal-Wallis with Dunn’s post-hoc test with Bonferroni correction. Non-significant comparisons are omitted from all plots.

### Neighborhood context influences immune cell activation marker profile in MM

We next analyzed functional marker expression profiles of immune cells within tumor and immune NBHs to gain further insights into their activation. DCs and HLA-DR^+^ macrophages (mascs)/ monocytes (monos) from ADAPTIVE_IMMUNE had the highest expression of both HLA-DR and IDO when compared with the HIF1A_IMMUNE and tumor NBHs, which could reflect a more vigorous DC/macrophage activation (Fig. 6E and F). GZMK^+^ T cells have cytotoxic properties and are often associated with a memory-like T cell phenotype.^30^ We found GZMK to be higher expressed in ADAPTIVE_IMMUNE and HIF1A_IMMUNE in CD8^+^ T cells when compared with T cells from PC_MYELOID and PC_OXPHOS (Fig. 6G). Taken together, the expression of HLA-DR, IDO, and GZMK in antigen-presenting cells (APCs) and CD8⁺ T cells may indicate increased activation and an ongoing immune response within immune-enriched NBHs. In contrast, this expression was less prominent when the same cell types were located within tumor NBHs, possibly reflecting a more immunosuppressive microenvironment. However, we lack the necessary markers to further determine whether this immune response is immunostimulatory and cytotoxic or instead reflects a tolerogenic and exhausted phenotype.

### Neighbor preference between CD4^+^ T cell and PC provides prognostic information in MM

To investigate whether interactions between cells or neighborhoods carry prognostic value, we analyzed spatial features in relation to progression-free survival (PFS) within our dataset. Due to limited follow-up time, data on overall survival (OS) was not available. We stratified MM patients with disease progression data (N=63) into short (N=29, PFS<2 years) and long (N=34) PFS groups (Supplementary Table 4). Both groups displayed similar cellular abundances and neighborhood frequencies suggesting that abundance per se is not associated with disease progression (Fig. 7A, Supplementary Fig. 8A and B).

**Fig. 7.**
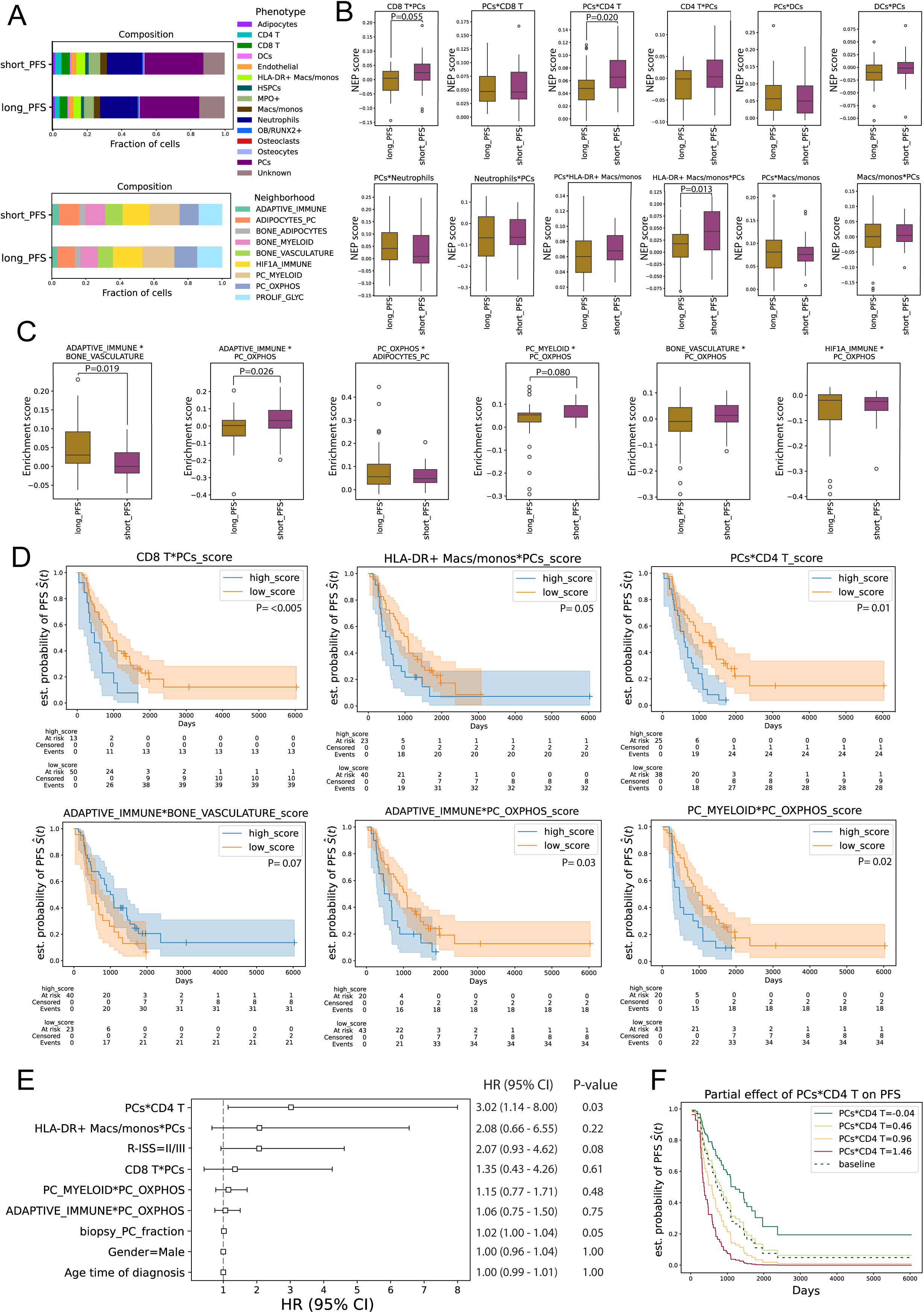
Neighbor preference between CD4^+^ T cell and PC provides prognostic information in MM. Patients were grouped based on their progression-free survival (PFS) into short_PFS (<2 years) (N=29) and long_PFS (>2 years) (N=34). **(A)** Cumulated cell type (top)- and neighborhood (bottom) abundances in groups of patients with short and long PFS. **(B)** Pairwise neighbor preference (NEP) scores between PCs and different immune cells comparing patients with short and long PFS. NEP scores are normalized per image. The median in each cohort is indicated by a horizontal solid line, whiskers extending to the most extreme data points within 1.5 times the interquartile range from the quartiles. Outliers are shown as individual points. P-values determined by Mann-Whitney-U-Test. Non-significant comparisons are omitted from all plots. **(C)** CellCharter-derived neighborhood enrichment scores between selected neighborhoods comparing patients with short and long PFS. The median in each cohort is indicated by a horizontal solid line, whiskers extending to the most extreme data points within 1.5 times the interquartile range from the quartiles. Outliers are shown as individual points. Outliers are shown as individual points. P-values were determined by Mann-Whitney-U-Test. Non-significant comparisons are omitted from all plots **(D)** Univariate Kaplan-Meier analyses of continuous PFS data and neighbor preference scores or neighborhood enrichment scores in the MM patient group (N=63). Statistical significance was assessed via log-rank test (p<0.05). Shaded areas indicate 95% confidence intervals. **(E)** Cox proportional hazard (CoxPH) model with elastic net regularization fitted using the significant univariate predictors from (D) and the clinical covariates age, gender, biopsy PC fraction and R-ISS score for 49 MM patients. Scores from (D) were scaled by factor 10 to ensure realistic visualization and interpretation in the CoxPH model. P-values were obtained by Wald test. **(F)** Predicted survival curves for the cohort using different values of the PC*CD4 T cell NEP score while controlling for other variables.

Given that immune–cancer cell interactions and immune exclusion have been linked to survival in other cancer types ^31,32^, we next investigated whether spatial patterns were associated with PFS in our dataset. Interestingly, neighbor preference of PCs and CD4^+^ T cells, and of HLA-DR^+^ macs/monos and PC differed significantly between the PFS groups, consistently showing increased preference scores in patients with short PFS (Fig. 7B). A similar trend was observed for neighbor preference of PCs and CD8^+^ T cells (Fig. 7B). To identify interactions on neighborhood level, we selected promising candidates after running a differential neighborhood enrichment analysis (Supplementary Fig. 8C). The two identified cancer NBHs PC_MYELOID and PC_OXPHOS as well as PC_OXPHOS and ADAPATIVE_IMMUNE showed increased enrichment in the short PFS group (Fig. 7C). In contrast, high enrichment scores between ADAPTIVE_IMMUNE and BONE_VASCULATURE were associated with the long PFS group (Fig. 7C).

To assess the prognostic relevance of neighbor preference scores and neighborhood enrichments identified in group comparisons, we evaluated their association with continuous PFS using univariate Kaplan–Meier analyses, applying optimal cutoff stratification for each score. Notably, stronger interactions between PCs and CD8⁺ T cells, CD4⁺ T cells, or HLA-DR⁺ macrophages/monocytes were each associated with shorter PFS (Fig. 7D). At the neighborhood level, enrichments between PC_MYELOID and PC_OXPHOS, as well as between ADAPTIVE_IMMUNE and PC_OXPHOS, were also significantly associated with shorter PFS in univariate analyses (Fig. 7D). Importantly, these associations with PFS were observed both at the cellular level (PC–T cell interactions) and at the neighborhood level (PC_OXPHOS–ADAPTIVE_IMMUNE proximity), suggesting a consistent biological phenomenon captured across spatial scales and through two independent analytical approaches.

Significant univariate predictors were subsequently analyzed in a Cox proportional-hazards (CoxPH) model including the clinical covariates age, gender, fraction of PCs in the biopsy and R-ISS staging, which was available for 49 patients (Fig. 7E). When including all previously identified univariate PFS predictors, only the neighbor preference score between PCs and CD4^+^ T cells remained as a variable significantly associated with increased risk of progression (Fig. 7E). The PCs*CD4^+^ T cell preference score also retained predictive value in a CoxPH model including all 63 patients – where only age, gender and fraction of PCs in the biopsy were available (Supplementary Fig. 8D). The predictive value of the PC*CD4⁺ T cell neighbor preference score is illustrated in a partial effects plot of predicted survival curves, showing that, after adjusting for other variables, higher scores are associated with an increased risk of progression, while lower scores correspond to a more favorable prognosis (Fig. 7F).

## Discussion

Due to the intricate nature of the TME in MM, spatial mapping of cellular interactions is crucial for understanding disease progression. In this study, we leveraged high-dimensional IMC to spatially resolve the BM microenvironment in MM at single-cell resolution. We identified plasma cell niches characterized by distinct metabolic traits, as well as variations in the local microenvironment surrounding bone across different patient groups. Moreover, we demonstrate that spatial interactions between PC and immune cells can provide prognostic information.

The bone disease (BD) of MM patients affects about 80% of patients at diagnosis and represents a severe clinical problem. Several factors secreted from malignant plasma cells (PCs) or other cells in the TME can promote osteoclast activity, or inhibit osteoblast function in MM.^18,19^ A central finding from this study is that patients with BD have an increased abundance of plasma cells in the vicinity of osteoclasts compared with MM patients without BD, even though the total number of MM cells did not differ between the groups. Thus, direct contact between MM cells and osteoclasts is a key feature distinguishing patients with BD from those without, supporting previous findings.^21^ We also identified a bone-proximal metabolic phenotype of PCs in patients with BD. In BD patients, plasma cells near bone surfaces exhibited elevated expression of IL-32 and HIF1a. IL-32 is a non-conventional cytokine, regulated by HIF1a, which promotes osteoclastogenesis and bone loss.^22,33^ Interestingly, IL-32 is secreted in vesicles, indicating a primarily local effect – making its elevated expression near bone surfaces in BD patients particularly noteworthy.^22^ This distance-dependent signature was not observed in patients without BD, suggesting that either the bone niche itself differs between patients with and without BD, or that PCs respond differently to the same niche in the two groups. Notably, these PC phenotypes were not captured by global or non-spatial metrics, highlighting the value of spatially resolved analysis in uncovering pathophysiological mechanisms of MM. However, due to the limited sample size of patients without BD, these findings require further research.

Beyond single cells, we identified distinct BM NBHs that reflect both cellular composition and metabolic states. Two PC-dominated NBHs -PC_OXPHOS and PC_MYELOID-emerged with differences in growth patterns, vascularization, and metabolic phenotype. PC_OXPHOS was characterized by dense PC aggregates, enrichment of endothelial cells, and high expression of OXPHOS-related markers, indicating a vascularized and metabolically active niche. The existence of focal lesions containing sheets of malignant PCs is a well-described phenomenon in MM, both at the macro level^27^, and recently, at the micro level^11,12^. On a macro scale, Fluorine-18 fluorodeoxyglucose positron emission tomography with CT (18F-FDG PET/CT) is used to detect focal lesions in myeloma patients by measuring glucose consumption, which has led to the assumption that tumor cells primarily rely on glucose metabolism.^34,35^ However, our analyses reveal that PCs from focal clusters at the micro scale do not solely rely on glycolysis, as we observe a distinct OXPHOS-related phenotype in this NBH. In contrast, PC_MYELOID features dispersed PC growth, enrichment of myeloid immune cells, and elevated expression of glycolytic markers. These findings underscore that there is spatial and metabolic heterogeneity even within small regions of the MM BM. The distinct profiles in different regions of the BM suggest that malignant PCs vary in their individual properties such as activity, dormancy and other cellular processes. Supporting this notion, we observed a general trend towards lower proliferative activity in PCs located close to the bone surface, consistent with previous mouse studies demonstrating that niches of PC dormancy exist in the MM BM.^16,17^

We also found that tumor growth patterns influenced immune engagement. Larger PC aggregates in the PC_OXPHOS neighborhood were associated with reduced immune cell interaction, suggesting immune exclusion in metabolically active, vascularized niches. A similar phenomenon of T cell exclusion has been recently described for MM focal areas, where T cells were excluded from the tumor boarders in the absence of CLEC9A^+^ type 1 dendritic cells^11^, which play a crucial role in antitumor immunity.^36^ Importantly, we observed that interactions between immune cells and PCs bear prognostic significance. Interestingly, and contrary to expectations, increased interactions between immune cells and plasma cells were associated with shorter PFS. Notably, this negative association was consistently observed at both the cellular and neighborhood levels, as determined by two independent approaches. In particular, increased neighbor preference of CD4^+^ T cells to PCs was significantly associated with shorter PFS and remained significantly associated with increased risk for progression in a multivariate analysis including clinical parameters and grading system R-ISS.^5^ Previous studies linked increased CD4^+^ T cell density or frequencies to reduced PFS.^11,37^ However, these studies did not investigate direct interactions between PCs and CD4^+^ T cells. BM CD4^+^ Th cells promote MM cell proliferation in vitro^38^, which may provide a biological explanation for our finding. It may also be that the interacting CD4^+^ T cells could be regulatory cells (Tregs).^39^ Tregs foster an immunosuppressive microenvironment and directly promote PC survival within the BM ^39,40^. We did not include Treg markers in our antibody panel and were therefore not able to distinguish between Tregs and Th cells in this dataset. This limitation should be addressed in future studies.

In conclusion, our study provides a comprehensive spatially resolved map of the bone marrow microenvironment in MM and its precursor states, integrating high-dimensional imaging with clinical annotations across disease stages. By applying two independent, multi-resolution spatial analysis frameworks, we identify reproducible features, such as adaptive immune–plasma cell interactions and bone-distance–dependent metabolic profiles, that are associated with disease progression and bone involvement. Our work offers a valuable resource for the community and lays a strong foundation for incorporating spatial profiling into future risk stratification and therapeutic decision-making frameworks. Further studies in larger, prospectively followed cohorts with expanded immune phenotyping will be critical to validate and build upon these insights.

## Methods

### Patients

Formalin-fixed paraffin embedded (FFPE) iliac crest trephine bone marrow biopsies obtained at diagnosis from myeloma patients (N=65) and patients diagnosed with SMM (N=6) or MGUS (N=5) were obtained from Biobank1 (St.Olavs Hospital, Trondheim, Norway). 55 of MM patients were considered to have BD at diagnosis based upon the presence of one or more osteolytic lesion evident by low dose CT and/or fracture. 10 patients presented without BD, of those 6 developed lesions upon follow up (median time from noBD to BD: 1315 days). The median age at time of diagnosis for the MM patients was 65 years (34-83), with a gender distribution of 41 males (63.1%) and 24 females (36.9%). For the PFS analyses two patients from the MM cohort were excluded due to death of unknown reason before disease progression. The patient characteristics are summarized in Supplementary Table 1. All samples were acquired after giving consent. The study was approved by the regional ethics committee (REK 247909).

### Tissue preprocessing

The standard procedures for FFPE embedding include formalin fixation for a minimum of 24 hrs. before decalcification with EDTA for 6-18 hrs. Biopsies were then washed thoroughly and placed back on formalin, before dehydration and clearance using ethanol and Tissue-Clear before embedding in Histowax. For image acquisition, two regions of interest (ROIs) of 1 mm^2^ (1000×1000 pixels) were selected per patient. The two ROIs were selected to capture heterogeneity in the bone marrow tissue within each patient. Furthermore, as osteoclasts are present in low numbers in the bone marrow, the selection of ROIs were also based on Tartrate Resistant Acid Phosphatase (TRAP)-staining (Invitrogen, clone 26E5) of neighbor sections using IHC.

### HES and IHC staining procedures

4 µM sections of FFPE BM and tonsil biopsies were cut using a microtome and mounted on Superfrost+ slides. Paraffin was removed using Tissue-Clear, before rehydrated using descending grades of ethanol to water. For HES staining, sections were stained in Hematoxylin (Sigma-Aldrich, C.I.75290) followed by staining with Erythrosine (Sigma-Aldrich, 720-0179) in an automated slide stainer (Sakura Tissue-Tek (C) Prisma^TM^). Sections were rinsed in water and dehydrated in ascending grades of ethanol and stained with Saffron (Waldeck, 5A – 394,). Sections were rinsed in absolute ethanol and cleared in TissueClear before cover slipping (Sakura Tissue-Tek Glas). For IHC staining of bone marrow and tonsil, antigen retrieval was conducted using Tris-EDTA (pH9) at 97° C for 20 min using PT Link Pre-Treatment Module for Tissue Specimen (Dako). Sections were incubated with primary antibodies for 40 min with dilutions as specified in Supplementary Table 2, in Dako Cytomation Autostainer Plus (Dako). Dako REAL^TM^EnVision^TM^ detection system was used for visualization.

### Panel design and heavy metal conjugation of antibodies

Antibodies were purchased from standard Biotools (STB) when available (Supplementary Table 2). For antibodies not commercially available, antibodies were conjugated with heavy metal isotopes in-house, using the Maxpar Ab labeling kits (STB) according to manufacturer’s instructions (Supplementary Table 2). The expected abundance and expression levels of the different protein markers were evaluated when allocating and conjugating antibodies to different metal tags.

### Antibody validation and titration

Prior to conjugation, in-house conjugated antibodies were validated by IHC staining of both bone marrow biopsies and in some cases, tonsil (Supplementary Fig. 9). The specificity and distribution of expression was evaluated together with a pathologist. In-house conjugated antibodies not validated by IHC have been validated in previous publications with comprehensive information regarding antibody specificity and binding^13,41^. All antibodies were titrated and tested using IMC to ensure appropriate staining intensities. Furthermore, all antibodies were evaluated by inspecting IMC images to verify expected protein expression within different cell types.

### Staining for Imaging Mass Cytometry and image acquisition

To minimize batch effects, stocks of concentrated antibody cocktails were prepared for all samples and frozen at -80 °C. Immediately prior to antibody incubation, vials with antibody cocktails were thawed and diluted with BSA (Sigma-Aldrich, A7906) and PBS (Sigma-Aldrich, D8537) to a final concentration of 0.5% BSA. 4 µm tissue section on superfrost+ slides were incubated for 1-2 hrs. in a 60 °C incubator for paraffin removal. Sections were placed in UltraClear (J.T.Baker, 3905.2500PE) for tissue clearing, before rehydrated using descending grades of ethanol, before rinsed in distilled H_2_O. Antigen retrieval was conducted using Antigen Retrieval Reagent Basic (Agilent, S236784-2), diluted 1:10 with dH_2_O at 97 °C for 30 min using PT Link Pre-Treatment Module for Tissue Specimen (Dako). Next, sections were washed in EnVision Flex Wash Buffer (20X) (Dako, K800721-2) and subsequently encircled with a PAP pen before slides were placed in a hydration chamber, and incubated with Superblock (Thermo Fisher, 37580) for 45 min at RT. Next, sections were incubated with diluted antibody cocktails overnight at 4 °C. Slides were then rinsed in EnVision Flex Wash buffer and PBS. Sections were further incubated with Cell-ID^TM^ Intercalator (62.5 nM)(201192A, STB) diluted in PBS for 30 min in a hydration chamber. Lastly, sections were rinsed with PBS and dH_2_O before airdried in RT. IMC was performed using Hyperion Imaging System (STB) coupled with a Helios Mass Cytometer (STB) with laser-ablation at a rastered pattern of 200 Hz. 2 ROIs of 1 mm^2^ were acquired from each patient, with a resolution of 1 µm/pixel.

### Data analysis

#### Data Preprocessing

Raw mcd-files were processed using the dockerized version of the IMC processing toolkit Steinbock.^42^ In brief, images were preprocessed with a hot pixel filter value of 50. Whole-cell segmentation was performed with DeepCells’ MESMER^43^, which has already been successfully applied in multiplex bone-marrow datasets^44,45^. HistoneH3, 191Ir and 193Ir were used as a combined nuclear channel and CD98, CD3, CD138 and CD45 as a combined membrane channel. Single-cell features were extracted using Steinbock’s object measurement module. Cells with an area smaller than 4 pixels were filtered out and each marker was censored to the 99^th^ percentile and subsequently arcsinh-transformed with cofactor 1. For downstream analysis, each cell in each sample was normalized using muon’s centered log ratio (CLR) for proteomics^46^ based on the total protein abundance per cell and then combined, as also described before^44^.

#### Bone labeling and calculation of distance to bone

Due to challenges related to bone detachment during antigen retrieval and IMC staining, we performed manual annotation of all bone areas using polygons in QuPath 0.5.1.^47^ For each ROI, the bone annotation was based on the HE-stained neighboring section, where bone areas are usually preserved. Bone masks were generated from polygon-derived coordinates. These masks were then used to compute the minimum distance to the nearest bone surface for each cell (label) using the *’scipy.spatial.distance_transform_edt’*^48^ function.

#### Quality check

All samples underwent visual inspection. Artifact regions were annotated using polygons in Napari^49^ and cells within these annotated regions were excluded from single-cell quantification tables. Segmentation quality was systematically verified through visual assessment.

#### Phenotyping

To achieve a specific cell type labeling we took a manifold approach (Supplementary Fig. 2B). The single cell data was batch-corrected using scanorama’s integration with scanpy.^50,51^ We then generated a visually annotated ground truth of 223,655 cells from 33 images by using SCIMAP’s prior hierarchical knowledge driven approach^52^ according to the documentation. In brief, a hierarchical decision matrix was created based on prior knowledge and scimap-specific weighting of the decisions *pos*, *anypos*, *allpos*, *neg*, *anyneg* and *allneg*. For every image, a threshold was set for each cell type defining marker using *’scimap.pl.gatefinder’*. Cell types were then generated using *’scimap.tl.phenotype_cells’* for each image separately. OCYs were identified by cells defined by absence of other cell type defining markers and localization within bone (Supplementary Fig. 10). However, loss of bone tissue during processing was evident in some biopsies, potentially leading to an under-representation of OCYs, and possibly OB/RUNX2+ cells and OCs that are attached to bone surfaces. Due to difficulties with nucleus and membrane detection, adipocyte annotation is incomplete. We acknowledge the inherent challenges associated with this by not integrating adipocytes or OCY into major downstream analyses and results.

The combined annotated single-cell tables were then split into training, validation and testing and then used to train a tree-based XGBoost^53^ model. We performed hyperparameter tuning with Optuna^54^ using the validation dataset and a stratified K-Fold of 10. The fine-tuned model subsequently achieved an F1 score of 0.86 on the test dataset. The remaining images were then annotated using the trained XGBoost model. Unknown cell types residing inside the bone were annotated as Osteocytes, while the remaining unknown cells were subjected to FlowSOM clustering^55,56^ with standard parameters. Three clusters with high expression of CD138, CD38 and IRF4 were reassigned to Plasma Cells. Because of the complex shape of macrophages inside tumor aggregates, we further refined Macrophage/Monocyte and Plasma cell annotations. For this, we manually set thresholds for CD68, HLA-DR and IRF4 on cells from 29 images and trained an XGBoost model to predict marker positivity on all cells from the remaining images. If previously annotated Plasma cells were positive for CD68 but negative for IRF4, they were relabeled as Macrophages. We purposely used IRF4 for the refinement, as it is a nuclear marker and therefore suffers less from lateral bleed present in highly aggregated tissue areas. Based on the presence of HLA-DR, these Macrophages/Monocytes were sub-classified as HLA-DR^+^ Macrophages/Monocytes. Finally, annotations were visually assessed by overlaying images with the cell type labels (Fig. 1B).

### Marker-guided Neighborhood analysis

#### Neighborhood definition

Spatial neighborhoods were created using CellCharter^24^ with recommended settings for spatial proteomics. First, a CellCharter-optimized trVAE model^57^ was trained with standard parameters to account for batch-effects and subsequently used for low-dimensional embeddings. Next, a spatial neighbor graph based on delaunay triangulation was created using Squidpy^58^ and long links were removed using the 95^th^ percentile. Neighbour cell features were aggregated up to 3 cell layers using the trVAE embeddings. Before running spatial clustering, the optimal cluster size of 9 was determined using the cluster stability function *’cellcharter.tl.ClusterAutoK’*with *n_clusters=(4,20).* Subsequently, neighborhoods were defined using CellCharter’s spatial clustering with *n_clusters=9*.

#### Enrichments

For phenotype enrichments inside each neighborhood, we used *’cellcharter.gr.enrichment*’ with p values generated by 250 permutations. Values are presented as log2fc and represent the fold-enrichment of a cell type in a neighborhood compared to the average abundance.

Neighborhood enrichment scores were calculated using ‘cellharter.gr.nhood_enrichment’ with p values generated by 250 permutations. Positive scores represent increased, negative scores decreased co-occurrence compared to spatial randomness calculated by permutation.

#### Connected components stability

To assess the tendency of aggregation, we used *’cellcharter.gr.connected_components’,* which produces connected component labels if cells from the same neighborhood are connected on the spatial neighbor graph and a user-defined threshold of a minimum number of cells is reached. We systematically increased the threshold from 1 to 300. In each step, we then calculated the fraction of cells from each neighborhood that still reside in a connected component based on the spatial neighbor graph.

#### Analysis of aggregated neighborhood cells

For analysis performed on aggregated neighborhood cells, we used *’cellcharter.gr.connected_components’* with *min_cells=60* and continued analysis only with cells from each neighborhood that still resided in a connected component. For comparing markers among the same cell types inside the aggregates, we used the mean expression (CLR) per image to avoid single-cell-level comparisons, which would introduce pseudoreplication bias by treating thousands of non-independent cellular measurements as statistically independent observations. Following this, we quantified the mean distance to the next bone surface.

#### COZI co-localization analysis

To determine neighbor preference scores between cell types, we calculated permutation-based z-scores using COZI.^29^ By using COZI, we did not assume symmetry in cell-cell neighbor preferences but gained directional preference scores between cell type pairs. We defined the nearest neighbors of a cell with Delaunay triangulation and randomly permuted cell type labels 300 times per image. We normalized the z-scores of an image by the square root of the total cell count in that image. This approach was recommended by the authors when encountering large differences in cell counts between images. For assessing neighbor preference of plasma cells and immune cells, we grouped all immune cells (CD4 T, CD8 T, DCs, Neutrophils, Macs/monos, HLA-DR^+^ Macs/monos) into one ‘immune cells’ label and subsequently ran COZI as stated above.

#### Spatial cell abundances

To determine the abundances of specific cell types in the radius of a target cell, we used scimaps *’*spatial_count*’* function, which calculates the fraction of each cell type in a specific radius around the target cell type. To account for differences in the abundance of the target cell, the median per image was taken.

#### Progression-free survival analysis

Patients were grouped into long PFS (>2 years) (N=34) and short PFS (<2 years) (N=29). For patient-specific neighbor-preference scores we used the COZI results generated before and focused on PC-immune interactions. For each patient, we took the mean score of both ROI, which generally showed the same positive or negative orientation. Differential neighborhood enrichments between patient cohorts were analyzed using *’cellcharter.gr.diff_nhood_enrichment*’ with PFS cohort labels. To extract the patient-specific neighborhood enrichment scores, we ran *’cellcharter.gr.nhood_enrichment*’ for each patient separately on the previously constructed spatial neighbors graph. Univariate Kaplan-Meier analyses were performed using the lifelines^59^ packages. First, previously calculated scores were dichotomized into high- and low score categories. Kaplan-Meier survival curves were then generated using these stratified groups, modeling PFS durations (time-to-event) with corresponding censoring indicators (event status). Between-group survival distributions were statistically compared using the log-rank test. The optimal cohort-derived cutoff for each score was determined with Log-Rank Optimization.

Variables showing significant univariate associations (p<0.05) were combined with clinical covariates (age, gender, fraction of PCs in biopsy, R-ISS) in a Cox proportional hazards (CoxPH) regression model regularized through elastic net penalty (α = 0.5, λ = 0.01). Given the low range of enrichment and interaction scores, we multiplied each score by a factor of 10 prior to modeling. This rescaling ensures that the hazard ratios presented in the CoxPH plot correspond to realistic changes in the predictors, as a one-unit increase in the original scale would not be observed in our data. The proportional hazards assumption was verified through analysis of scaled Schoenfeld residuals. Every covariate exceeded α=0.05 and therefore no violations were present. Multivariable effect estimates were visualized via forest plots displaying log-transformed hazard ratios (log[HR]) with 95% confidence intervals.

#### Software and statistical testing

All analyses were carried out in Python (3.9, 3.10 or 3.11 depending on the virtual environment) (Table 1). If not indicated otherwise, Mann-Whitney-U- or Kruskal-Wallis-Test with Dunn’s post-hoc test with Bonferroni correction were performed for testing statistical significances.

**Table 1:** Main software packages without associated dependencies.

| Name | Version | Language | Link |
| --- | --- | --- | --- |
| Scimap | 2.0.5 | Python 3.10 | https://scimap.xyz/ |
| CellCharter | 0.3.2 | Python 3.9 | https://cellcharter.readthedocs.io/en/latest/ |
| Steinbock | 0.16.1 | docker/bash | https://bodenmillergroup.github.io/steinbock/latest/ |
| XGBoost | 2.1.0 | Python 3.10 | https://xgboost.readthedocs.io/en/release_3.0.0/ |
| Squidpy | 1.6.1 | Python 3.9 | https://squidpy.readthedocs.io/en/stable/ |
| Scanpy | 1.10.3 | Python 3.9 | https://scanpy.readthedocs.io/en/stable/ |
| scarches | 0.6.1 | Python 3.9 | https://docs.scarches.org/en/latest/ |
| Scikit-learn<sup>60</sup> | 1.5.2 | Python 3.9 | https://scikit-learn.org/ |
| Scikit-image<sup>61</sup> | 0.22.0 | Python 3.11 | https://scikit-image.org/ |
| scanorama | 1.7.4 | Python 3.10 | https://github.com/brianhie/scanorama |
| Optuna | 3.6.1 | Python 3.10 | https://optuna.readthedocs.io/en/stable/ |
| COZI (fork of scimap) | 0.1.0 | Python 3.10 | https://github.com/SchapiroLabor/scimap_COZI |
| lifelines | 0.30.0 | Python 3.10 | https://lifelines.readthedocs.io/en/latest/ |
| scipy | 1.13.1 | Python 3.9 | https://scipy.org/ |
| Scikit-posthocs<sup>62</sup> | 0.11.4 | Python 3.9 | https://scikit-posthocs.readthedocs.io/en/latest/ |
| muon | 0.1.6 | Python 3.9 | https://muon.readthedocs.io/en/latest/index.html |

All analysis scripts including additional information will be deposited on GitHub: https://github.com/SchapiroLabor/myeloma_imc

#### Additional Material

The data including images, segmentation masks and the single-cell table will be uploaded to a public data repository upon publication.

## Acknowledgements

We thank Heidi Ødegaard Notø, Idun Dale Rein, Monica Bostad and Ingunn Nervik for technical assistance. The immunohistochemistry was performed at the Cellular and Molecular Imaging Core Facility (CMIC) – Histology, Norwegian University of Science and Technology (NTNU). CMIC is funded by the Faculty of Medicine and Health Sciences at NTNU and Central Norway Regional Health Authority. The IMC image acquisition was performed at the Flow Cytometry Core Facility, Institute for Cancer Research, Radiumhospitalet, Oslo University Hospital (OUS). The project has been funded by the Regional Health Authorities in Central Norway (Samarbeidsorganet), The Norwegian Cancer Society (#198161), the joint Research Committee (#30437) and the Cancer fund at St. Olavs Hospital. The authors gratefully acknowledge the data storage service SDS@hd supported by the Ministry of Science, Research and the Arts Baden-Württemberg (MWK), support by the state of Baden-Württemberg through bwHPC and the German Research Foundation (DFG) through grant INST 35/1314-1 FUGG, INST 35/1503-1 FUGG and INST 35/1597-1 FUGG. We acknowledge the use of OpenAI’s ChatGPT / Gemini 2.5 in this project which assisted by adapting the language to improve clarity and coherence while ensuring that the original scientific content remained intact. L.H. is supported by the Clinician Scientist Program of the Medical Faculty of the University of Heidelberg. L.H., C.S. and D.S. are supported by the Bruno and Helene Jöster Stiftung. C.S. and D.S. are supported by the German Federal Ministry of Education and Research (BMBF 01ZZ2004). D.S. is supported by the Ministry for Science, Research and Science Baden-Württemberg “MULTI-SPACE”; the Multi-dimensionAI project (CZS-Project number: P2022-08-101) was made possible by funding from the Carl-Zeiss-Stiftung; the Ministry for Science, Research and Science Baden-Württemberg „AI Health Innovation Cluster” and research funding from Cellzome, a GSK company.

## Notes

**Conflict of interest disclosure**: D.S. reports funding from GSK and received fees/honoraria from Immunai, Noetik, Alpenglow and Lunaphore. T.S.S. has received honoraria for lectures and educational material: Takeda, Celgene, Amgen, Johnsen&Johnsen/Janssen-Cilag, Abbvie, Pfizer; Consultancy: Bristol Myers Squibb, GSK, Sanofi, Pfizer, Menarini Group; Advisory board consultancy: Amgen, Celgene, GSK, Johnsen&Johnsen/Janssen-Cilag, Sanofi, Bristol Myers Squibb. The remaining authors declare no competing interests.

### Competing Interest Statement

D.S. reports funding from GSK and received fees/honoraria from Immunai, Noetik, Alpenglow and Lunaphore. T.S.S. has received honoraria for lectures and educational material: Takeda, Celgene, Amgen, Johnsen&Johnsen/Janssen-Cilag, Abbvie, Pfizer; Consultancy: Bristol Myers Squibb, GSK, Sanofi, Pfizer, Menarini Group; Advisory board consultancy: Amgen, Celgene, GSK, Johnsen&Johnsen/Janssen-Cilag, Sanofi, Bristol Myers Squibb. The remaining authors declare no competing interests.

