## Supplementary material for "Highly multiplexed imaging recovers immune and metabolic niches in multiple myeloma associated with disease progression and bone involvement"

Supplementary Figure 1

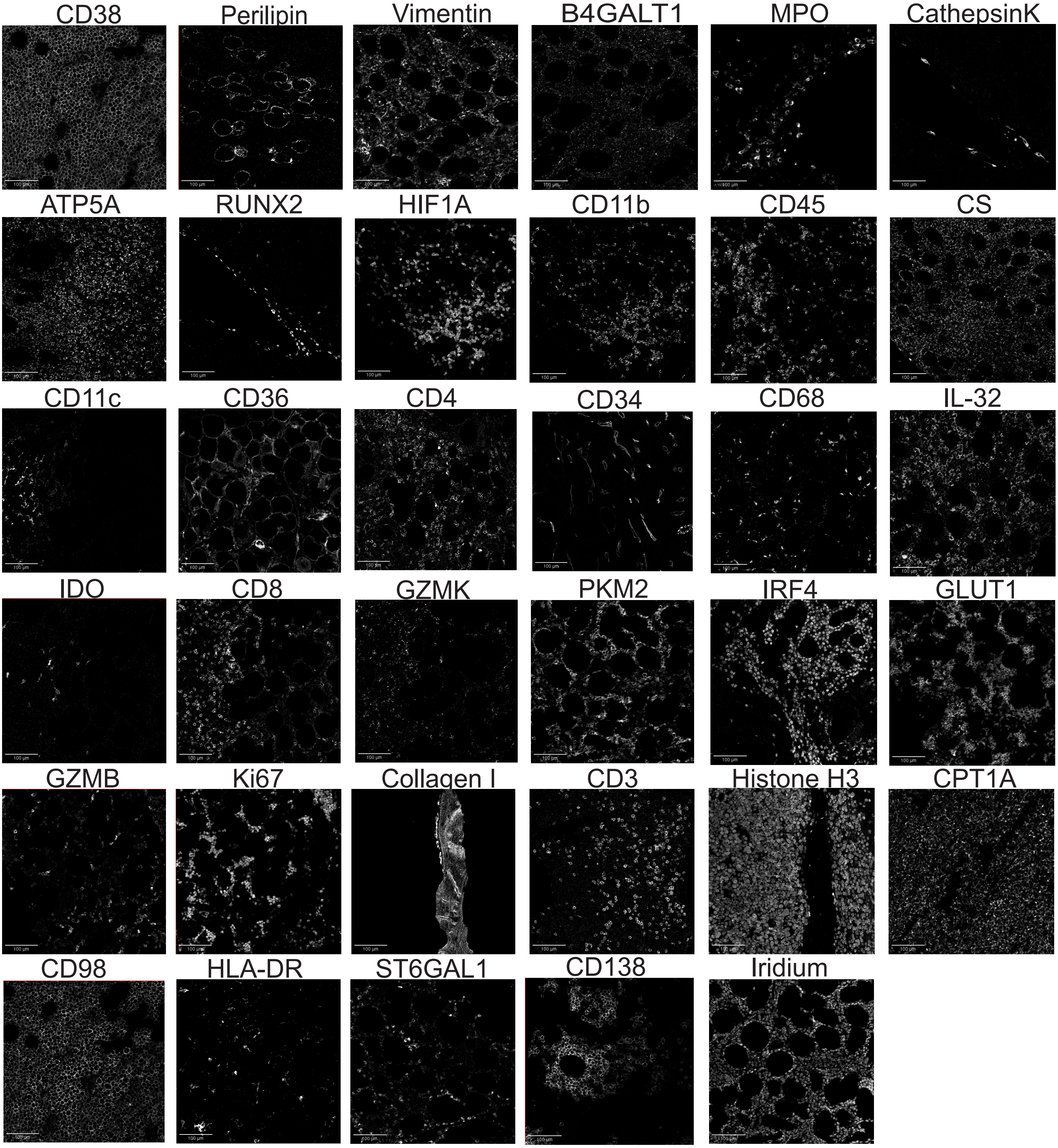

**Fig. S1. Visualization of individual marker signals from the 35-antibody IMC panel.**

Raw IMC images displaying a cropped region of representative ROIs and the individual signal from each antibody used in the study.

Supplementary Figure 2

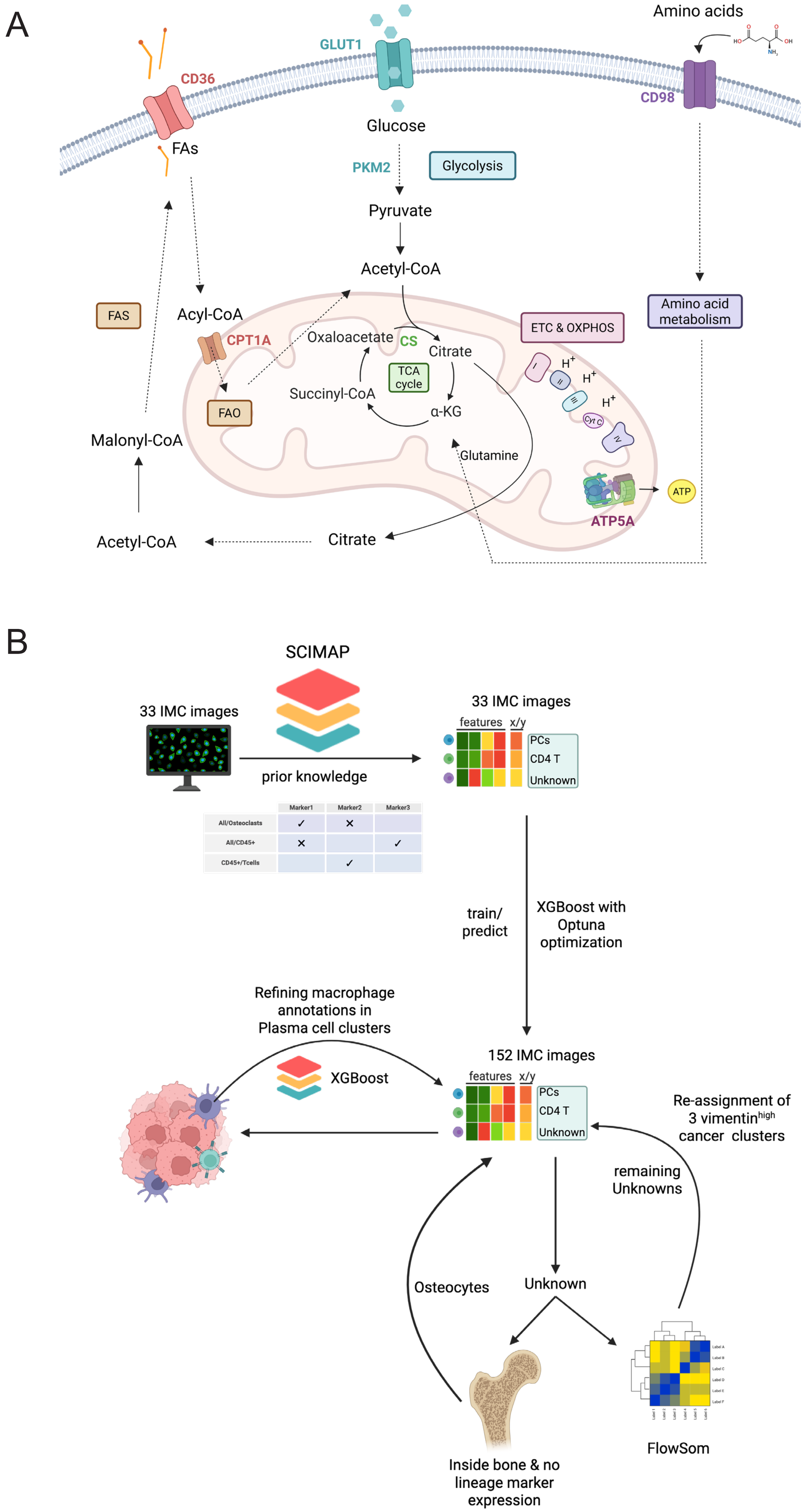

**Fig. S2. Schematic overview of pathways and metabolic markers included in IMC panel**

**(A**) Schematic overview of main metabolic pathways with highlighted markers included in the 35-antibody IMC panel used in the study. **(B)** 33 images were manually annotated using scimap's hierarchical prior knowledge method. Using these 33 images as a training/validation cohort, an optuna-optimized Xgboost model was built to predict annotations on all remaining images. All unknown cell types were clustered using FlowSom and the cancer clusters missed by XGBoost were re-assigned. Osteocytes were labeled from remaining unknown cells if located inside a bone mask and negative for all cell type-defining markers. Macs/monos annotation was refined according to the method section given their complex shape in cancer aggregates. Created with BioRender.com

### Supplementary Figure 3

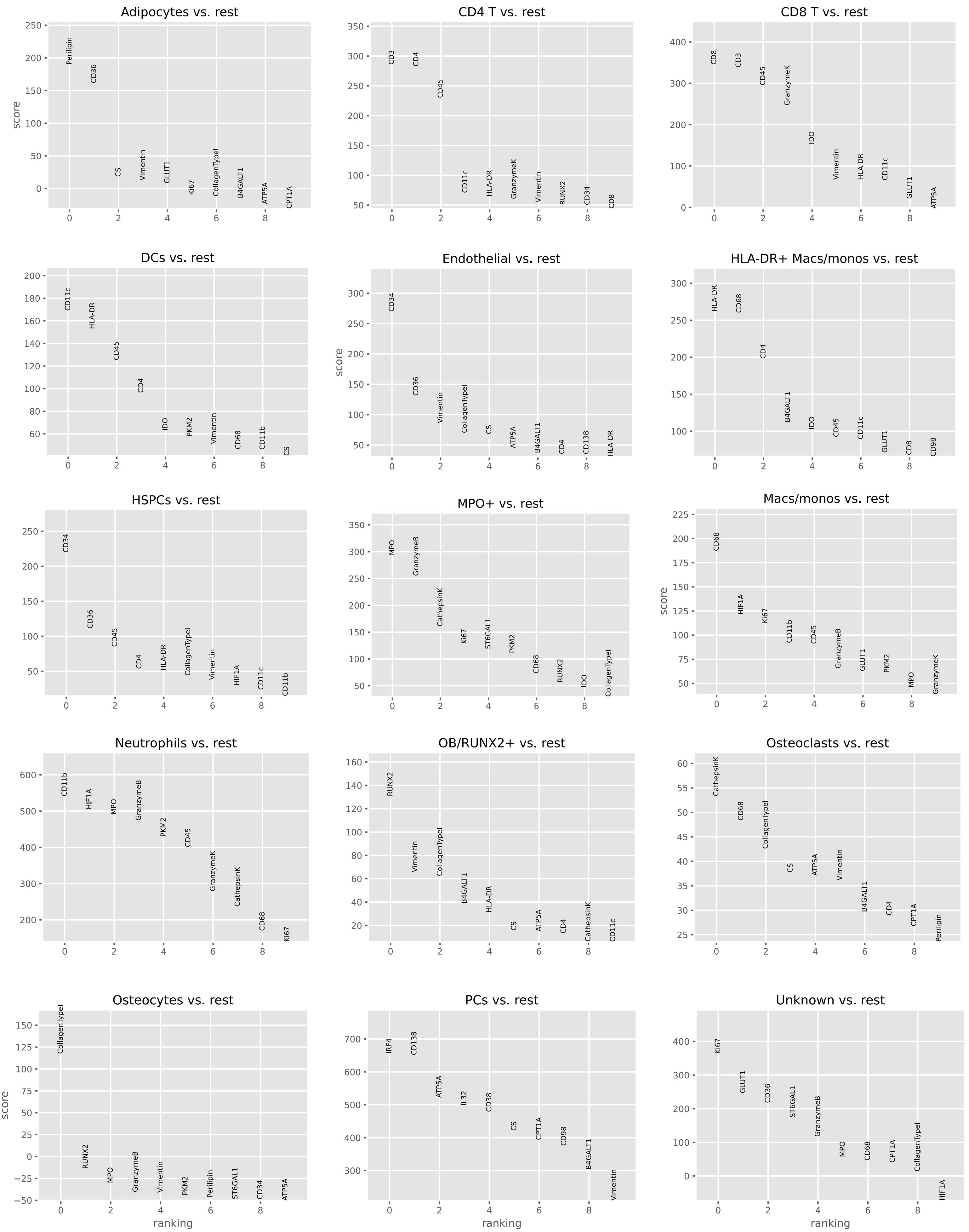

**Fig. S3. Top differentially expressed markers align with established cell type profiles**

Z-scores for the top 10 ranked genes for each cell type compared against all other cell types computed by scanpy's rank_genes_groups test with Wilcoxon's rank sum test.

Supplementary Figure 4

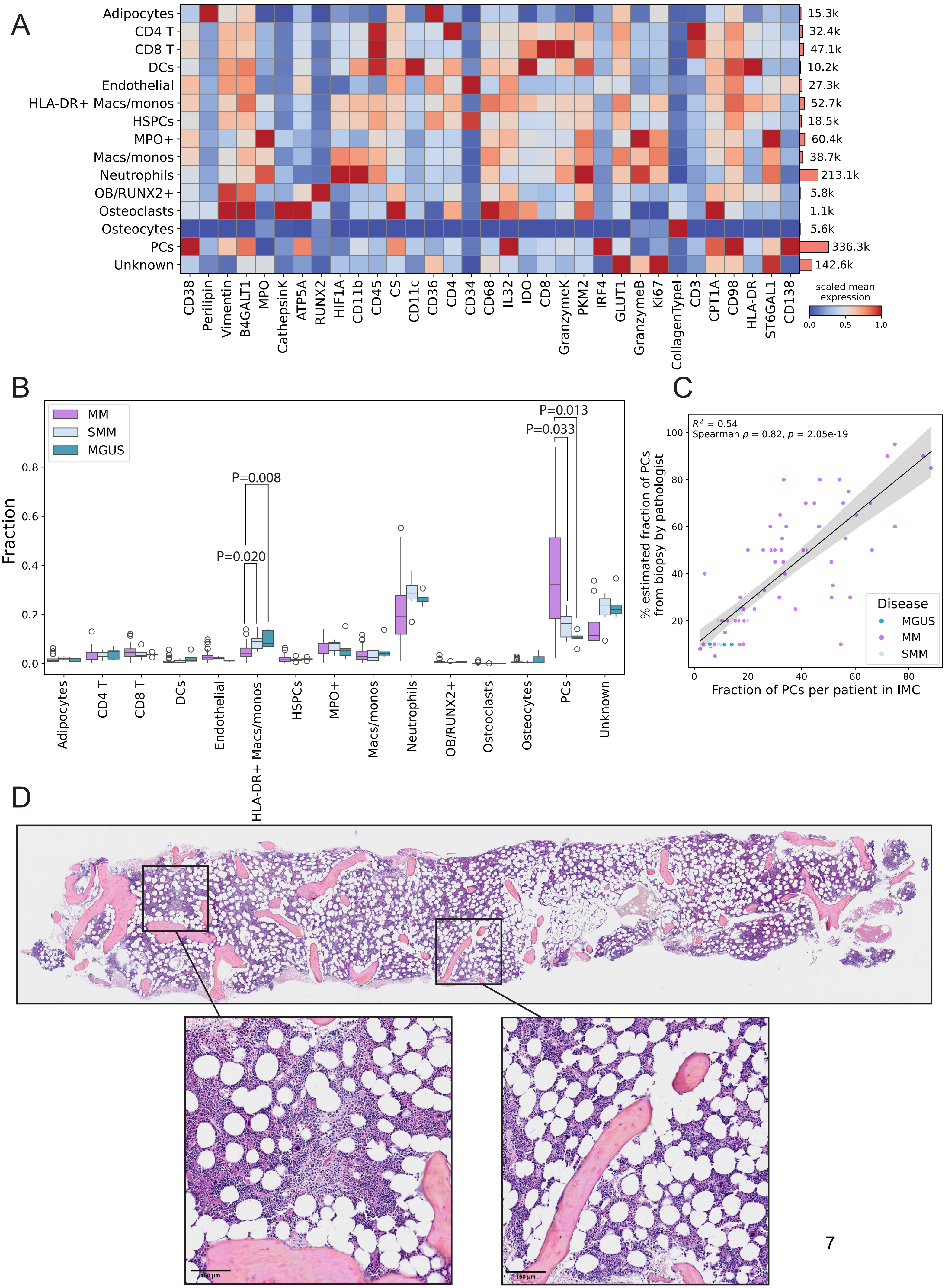

**Fig. S4. Cell typing aligns with common marker profiles and pathologist- estimated cancer burden**

**(A)** Heatmap showing arcsinh transformed scaled mean expression of all markers across all annotated cell types. **(B)** Cell type frequencies of all cell types in MM, SMM and MGUS patients. The median in each cohort is indicated by a horizontal solid line, whiskers extending to the most extreme data points within 1.5 times the interquartile range from the quartiles. Outliers are shown as individual points. P-values were determined by Kruskal-Wallis with Dunn's post-hoc test with Bonferroni correction. Non-significant comparisons are omitted from the plot. **(C)** Spearman correlation coefficients of the fraction of PCs per patient in the IMC dataset and the corresponding percentage of PCs estimated by the pathologists in the whole diagnostic biopsy. Spearman's $\rho = 0.82$, $p = 2.05*10^{-19}$ from correlation analysis; $R^2 = 0.54$ calculated from linear regression. **(D)** Example ROI selection from a complete HES-stained BM biopsy.

Supplementary Figure 5

A

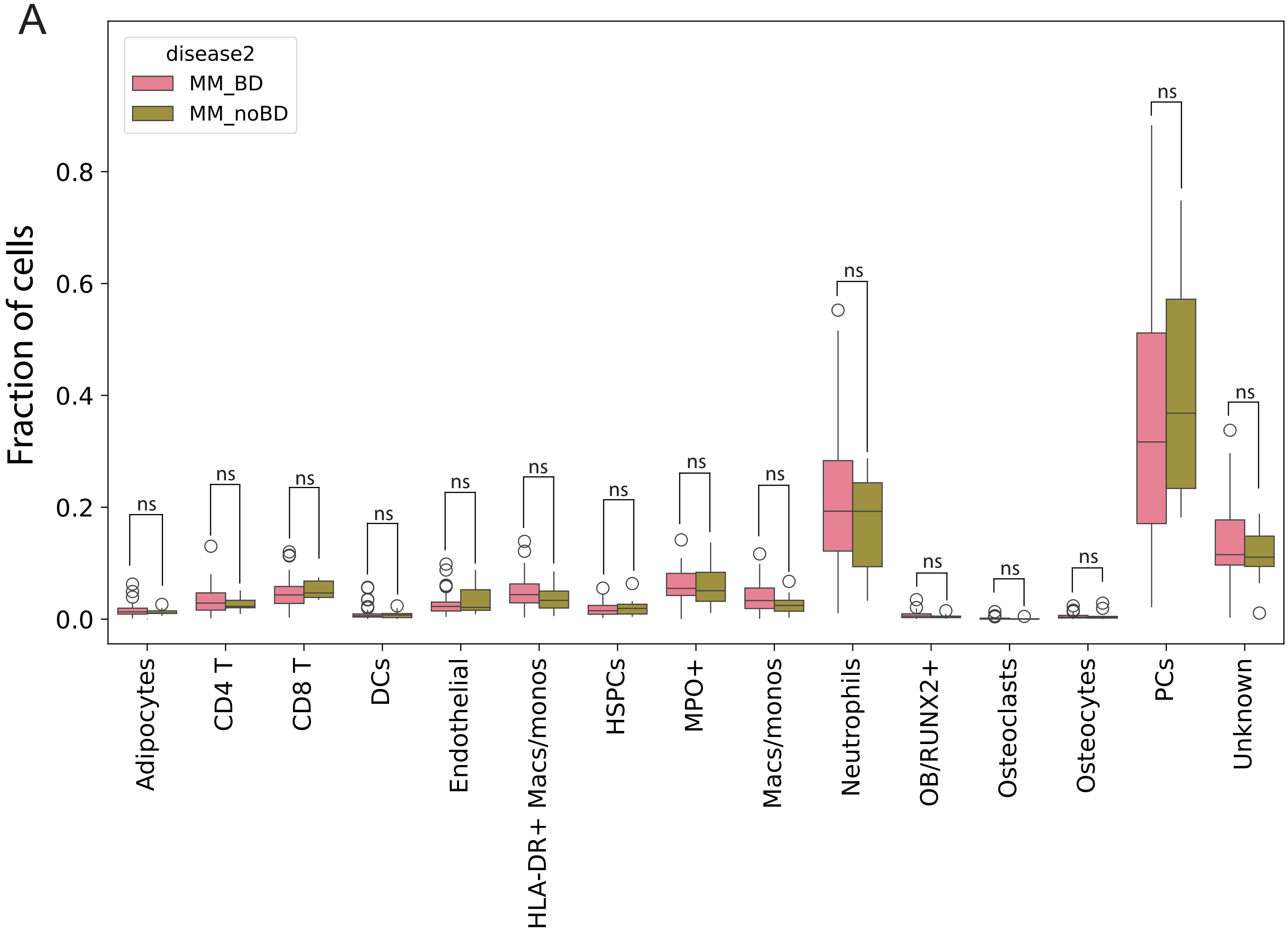


B

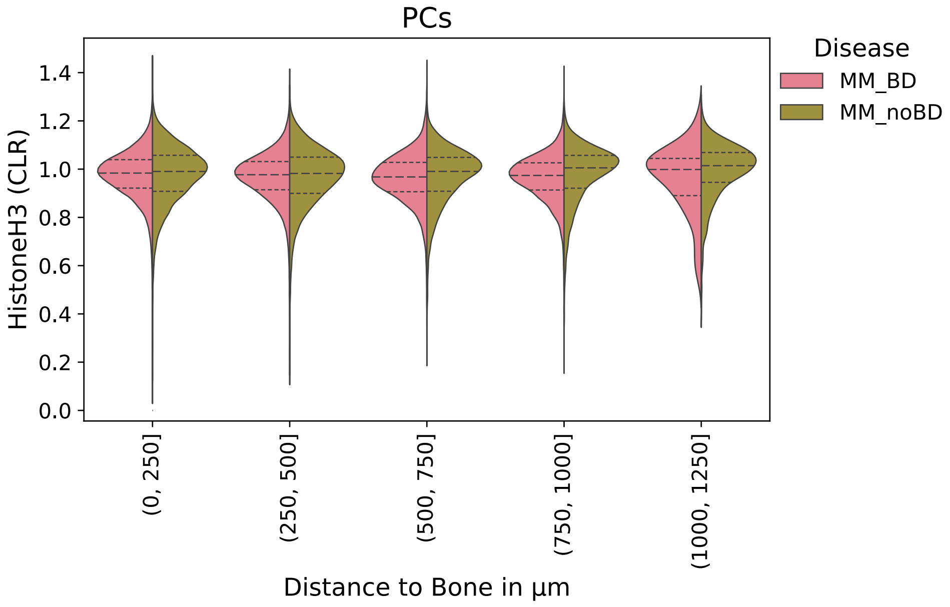

**Fig. S5. Cell type abundance is similar across BD and noBD patients**

**(A)** Cell type frequencies of all cell types between BD and noBD patients. The median in each cohort is indicated by a horizontal solid line, whiskers extending to the most extreme data points within 1.5 times the interquartile range from the quartiles. Outliers are shown as individual points. P-values determined by Mann-Whitney-U-Test. **(B)** CLR-normalized mean expression of HistoneH3 in PCs, stratified into five distance intervals based on distance to bone.

Supplementary Figure 6

A

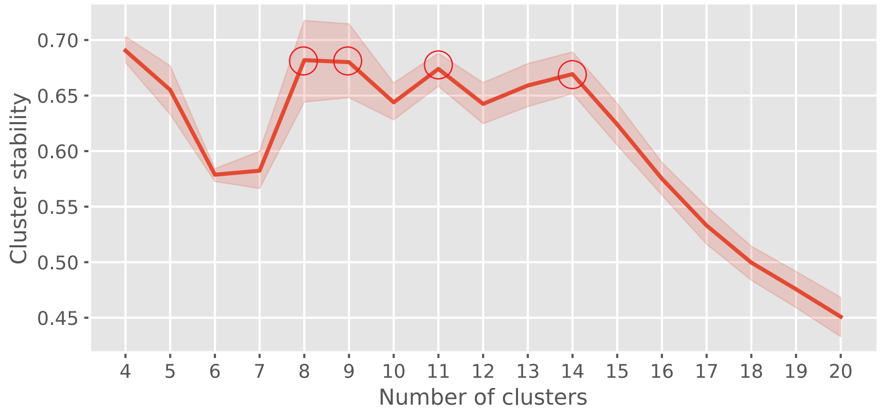


B

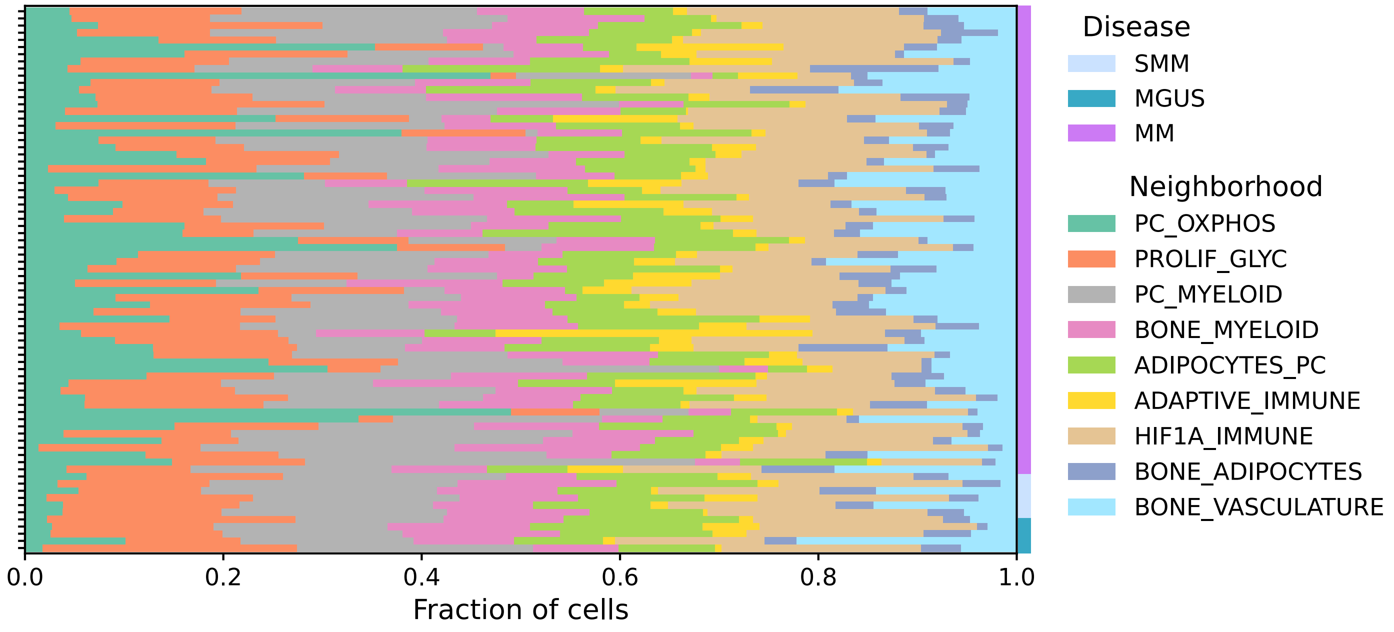

**Fig. S6. NBH cluster stability and patient-specific neighborhood distribution**

**(A)** Cluster stability (y-axis) quantified through average Fowlkes-Mallows Index (FMI) between consecutive cluster sizes (K±1) across 5 replicates. A cluster size of 9 was selected from stability peaks (circles) for downstream analyses. **(B)** Cumulated neighborhood composition in individual MM, SMM and MGUS patients.

### Supplementary Figure 7

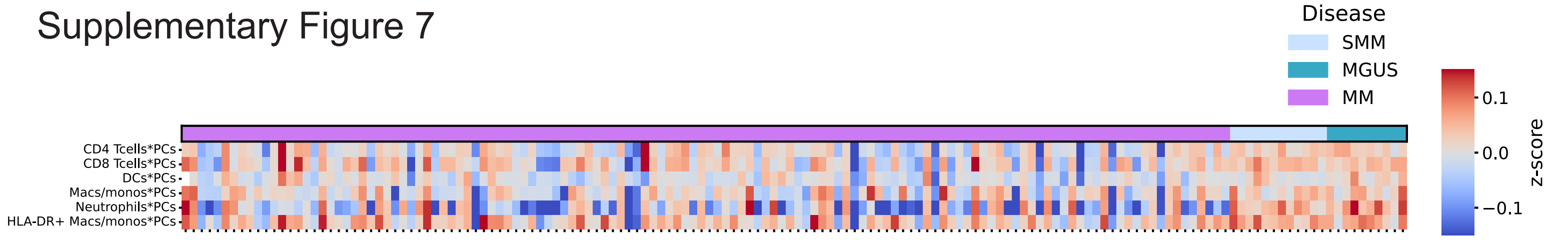

**Fig. S7. Distinct immune cell PC pair-wise neighbor preferences reveal disease-specific distributions**

Heatmap showing pairwise neighbor preference (NEP) scores between PCs and either CD4 T, CD8 T, DCs, Neutrophils, Macs/monos and HLA-DR+ Macs/monos. NEP scores are normalized per image.

Supplementary Figure 8

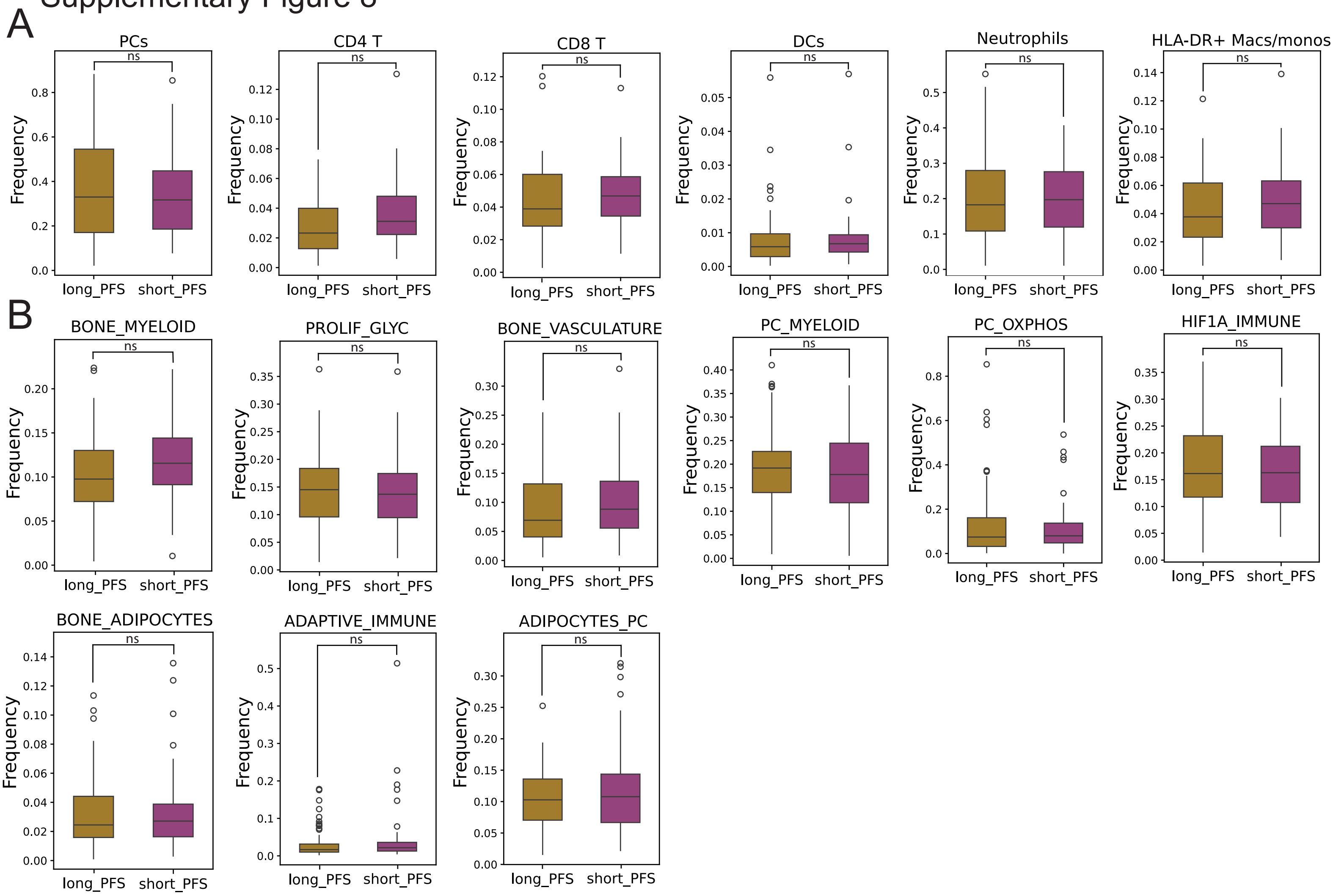


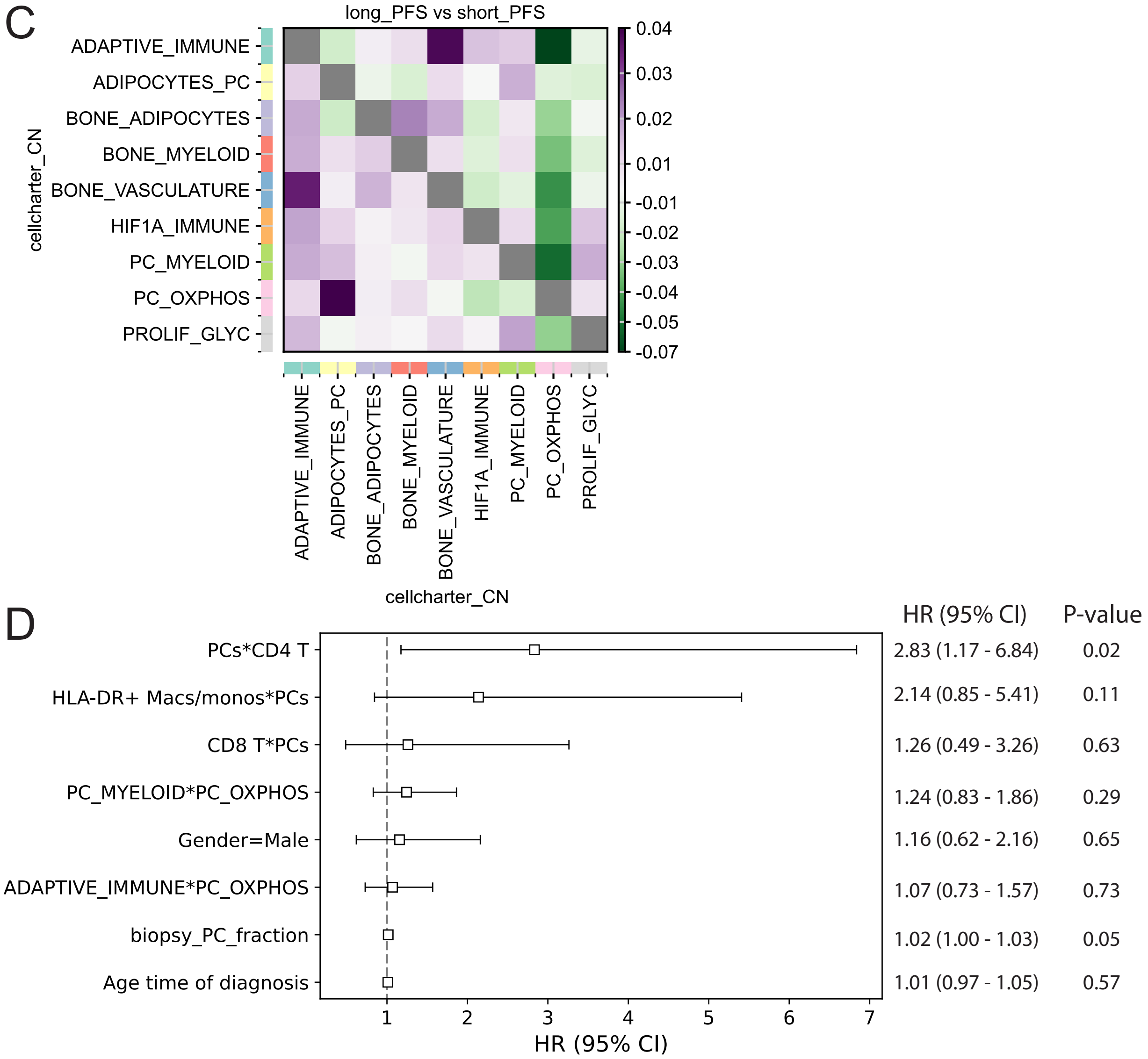

**Fig. S8. Cell type or neighborhood abundances do not distinguish PFS cohorts**

Cell type **(A)** and neighborhood **(B)** frequencies in short and long PFS patient groups. The median in each cohort is indicated by a horizontal solid line, whiskers extending to the most extreme data points within 1.5 times the interquartile range from the quartiles. Outliers are shown as individual points. The differences between groups were not significant (Mann-Whitney U test). **(C)** CellCharter's differential neighborhood enrichment applied to the two PFS patient groups, values represent the arithmetic difference between the enrichments scores of the two conditions. **(D)** Cox proportional hazard (CoxPH) model with elastic net regularization fitted using the significant univariate predictors from Figure 6D and the clinical covariates age, gender and biopsy PC fraction for all MM patients (N=63). Scores from Figure 6D were scaled by factor 10 to ensure realistic visualization and interpretation in the CoxPH model. P-values were obtained by Wald test.

Supplementary Figure 9

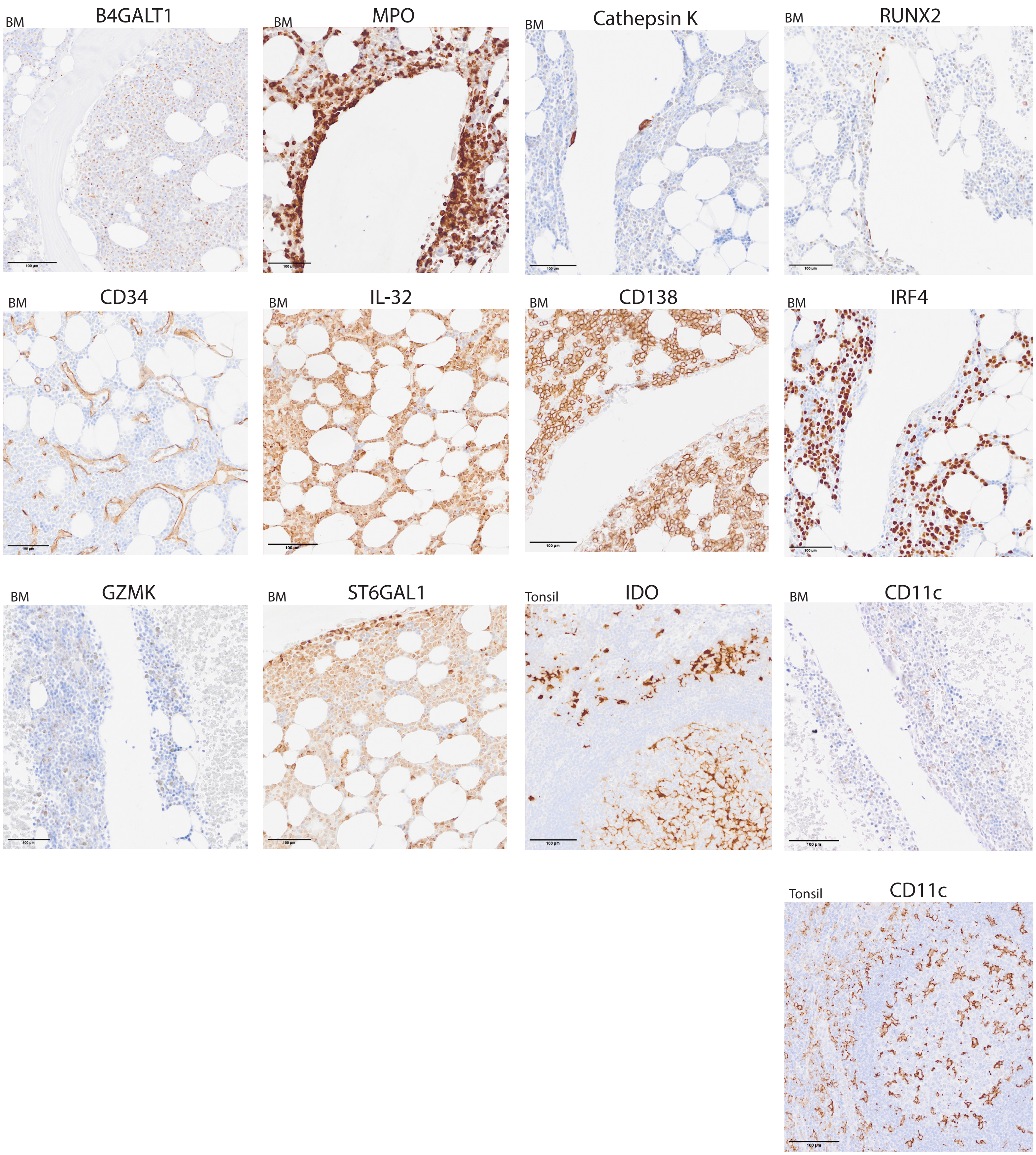

**Fig. S9. Immunohistochemical (IHC) validation of in-house conjugated antibodies.**

In-house conjugated antibodies were validated by IHC staining on BM tissue, or on tonsil tissue for antibodies against proteins with low abundance in BM (CD11c, IDO). Images display resulting staining pattern for each indicated antibody.

Supplementary Figure 10

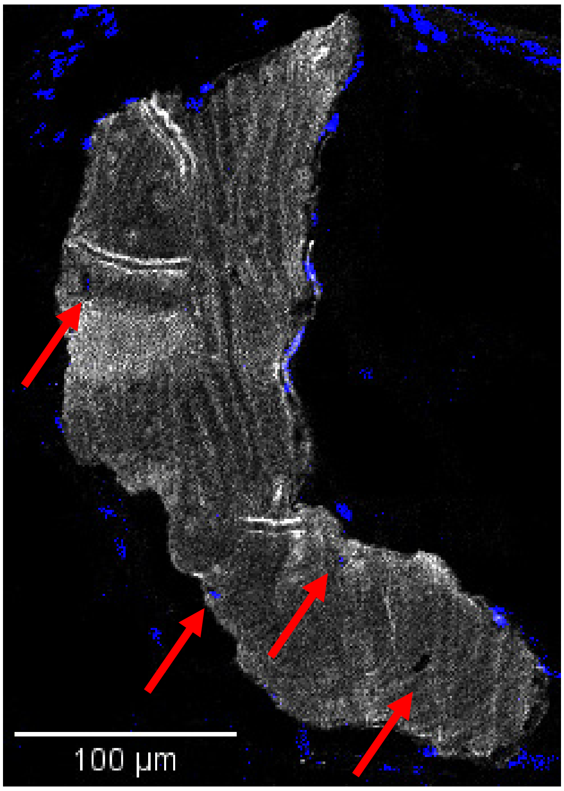

**Fig. S10. Schematic Overview of the cell annotation process.**

Raw IMC image visualized by Collagen I (white) and RUNX2 (blue), showing osteocytes inside bone (arrows).

Supplementary Table 1: Patient characteristics at diagnosis in untreated MM, SMM and MGUS patients

| Characteristics | MM (n=65) | SMM (n=6) | MGUS (n=5) |
|---|---|---|---|
| Age (median) y | 65 (34-83) | 71 (56-87) | 68 (51-75) |
| Sex (male) | 41 (63.07%) | 5 (83.3%) | 5 (100%) |
| Female | 24 | 1 | 0 |
| R-ISS I | 16 | 1 | 2 |
| R-ISS II | 30 | 3 | 1 |
| R-ISS III | 6 | 0 | 0 |
| R-ISS unknown | 13 | 2 | 2 |
| Bone disease upon diagnosis | 55/65 | 0/6 | 0/5 |
| ≥ 1 lesion | 4 | N/A | N/A |
| ≥ 3 lesion | 11 | N/A | N/A |
| ≥ 5 lesion | 18 | N/A | N/A |
| ≥ 10 lesion | 16 | N/A | N/A |
| Compression fractures | 24 | N/A | N/A |
| Fractures ossis longi/ossis plani | 19 | N/A | N/A |
| M-component (median) (g/dl) | 28.9 (0-69.8) | 13.75 (0.3-31.8) | 9 (5.9-17.9) |
| % PC from biopsy (median) | 45 % (5-95%)* | 22.5% (10-31%) | 9.9% (9.9-10%) |
| FISH t(11:14) | 6 | 1 | 0 |
| FISH t(14:11) | 1 | 0 | 0 |
| FISH t(4:14) | 12 | 0 | 0 |
| FISH del17p | 7 | 0 | 0 |
| FISH 1q21+ | 6 | 0 | 0 |
| FISH split IgH | 2 | 0 | 0 |
| FISH del13q | 1 | 0 | 0 |
| no translocations/deletions | 39[#] | 5[#] | 5[#] |
| LDH U/L (median) | 164 (71-441)[§] | 170 (133-202)[§] | 226 (143-271)[§] |
| B2-mikroglobulin mg/L (median) | 3.8 (1.7-24.9)[¤] | 3.3 (2-4.4)[¤] | 2.8 (2.2-3.9)[¤] |
| (short) PFS<2 years | 29 | N/A | N/A |
| (long) PFS>2 years | 36 | N/A | N/A |

* 1 patient <10%, 1 patient unknown

[#] 9 MM patients, 1 SMM and 1 MGUS patient without FISH data

[§] 11 MM patients unknown, 2 SMM patients unknown, 2 MGUS patients unknown

[¤] 13 MM patients unknown, 2 SMM patients unknown, 2 MGUS patients unknown

#### Supplementary Table 2: IMC antibody panel information

| Metal | Marker | Clone | Dilution | Cat.no | Supplier | Conjugated in-house | Validated by | IHC dilution |
|---|---|---|---|---|---|---|---|---|
| 141 Pr | CD38 | ERP4106 | 1:50 | 3141018D | Standard Biotools | | Standard Biotools | |
| 142Nd | Perilipin | Rabbit polyclonal, IgG | 1:50 | ab3526 | Abcam | yes | Aigular-Navarro, A.G. et.al. (2) | |
| 143 Nd | Vimentin | D21H3 | 1:1000 | 3143027D | Standard Biotools | | Standard Biotools | |
| 144 Nd | B4GALT1 | Rabbit polyclonal, IgG | 1:50 | HPA010807-100UL | Atlas antibodies | yes | IHC | 1:100 |
| 145 Nd | MPO | EPR20257 | 1:4000 | ab221847 | Abcam | yes | IHC | 1:1000 |
| 146Nd | Cathepsin K | EPR19992 | 1:600 | ab223140 | Abcam | yes | IHC | 1:1000 |
| 148Nd | ATP5A | 15H4C4 | 1:800 | ab14748 | Abcam | yes | Hartmann, F.J. et.al. (1) | |
| 149 Sm | RUNX2 | CL0232 | 1:200 | AMAb90591 | Atlas antibodies | yes | IHC | 1:300 |
| 150Nd | HIF1A | EP1215Y | 1:600 | ab210073 | Abcam | yes | Hartmann, F.J. et.al. (1) | |
| 151 Eu | CD11b | EPR1344 | 1:300 | ab209970 | Abcam | yes | Clone validated by Standard Biotools | |
| 152Sm | CD45 | D9M8I | 1:200 | 3152018D | Standard Biotools | | Standard Biotools | |
| 153Eu | CS | EPR8067 | 1:200 | ab233838 | Abcam | yes | Hartmann, F.J. et.al. (1) | |
| 154Sm | CD11c | EP1347Y | 1:200 | ab216655 | Abcam | yes | IHC | 1:600 |
| 155Gd | CD36 | D8L9T | 1:600 | 39914SF | Cell Signaling Technology | yes | Hartmann, F.J. et.al. (1) | |
| 156 Gd | CD4 | ERP6855 | 1:50 | 3156033D | Standard Biotools | | Standard Biotools | |
| 158Gd | CD34 | QBEND/10 | 1:400 | MA1-10202 | Invitrogen | yes | IHC | 1:50 |
| 159 Tb | CD68 | KP1 | 1:800 | 3159035D | Standard Biotools | | Standard Biotools | |
| 160 Gd | IL-32 | Rabbit polyclonal, IgG | 1:100 | ab37158 | Abcam | yes | IHC | 1:100 |
| 161 Dy | IDO | D5J4E | 1:50 | 91473 | Cell Signaling Technology | yes | IHC | 1:400 |
| 162 Dy | CD8a | C8/144B | 1:100 | 3162034D | Standard Biotools | | Standard Biotools | |
| 163 Dy | Granzyme K | EPR24601-164 | 1:100 | ab282714 | Abcam | yes | IHC | 1:1000 |
| 164Dy | PKM2 | D78A4 | 1:400 | 4053BF | Cell Signaling Technology | yes | Hartmann, F.J. et.al. (1) | |
| 165Ho | IRF4 | EP5699 | 1:100 | ab240071 | Abcam | yes | IHC | 1:250 |
| 166 Er | GLUT1 | EPR3915 | 1:1000 | ab196357 | Abcam | yes | Hartmann, F.J. et.al. (1) | |
| 167 Er | Granzyme B | EPR20129-217 | 1:10 000 | 3167021D | Standard Biotools | | Standard Biotools | |
| 168 Er | Ki67 | B56 | 1:200 | 3168022D | Standard Biotools | | Standard Biotools | |
| 169 Tm | Collagen type I | Goat polyclonal | 1:800 | 3169023D | Standard Biotools | | Standard Biotools | |
| 170 Er | CD3e | Polyclonal, C-terminal | 1:50 | 3170019D | Standard Biotools | | Standard Biotools | |
| 171 Yb | Histone H3 | D1H2 | 1:4000 | 3171022D | Standard Biotools | | Standard Biotools | |
| 172 Yb | CPT1A | 8F6AE9 | 1:400 | ab128568 | Abcam | yes | Hartmann et.al (1) | |
| 173 Yb | CD98 | Polyclonal | 1:100 | 3173018D | Standard Biotools | | Standard Biotools | |
| 174 Yb | HLA-DR | YE2/36 HLK | 1:400 | 3174023D | Standard Biotools | | Standard Biotools | |
| 175 Lu | ST6GAL1 | 036 | 1:400 | MA5-29582 | Invitrogen | yes | IHC | 1:200 |
| 176 Yb | CD138 | MI15 | 1:200 | 356502 | Biolegend | yes | IHC | 1:100 |
| 191Ir | Nuclear stain | N/A | 1:2000 | 201192A | Standard Biotools | | | |
| 193 Ir | Nuclear stain | N/A | 1:2000 | 201192A | Standard Biotools | | | |

Supplementary Table 3: Patient characteristics at diagnosis in untreated MM patients, divided into bone disease (BD) and no BD groups

| Characteristics | BD (n=55) | noBD (n=10) |
|---|---|---|
| Age (median) y | 65 (34-83) | 67.5 (47-79) |
| Sex (male) | 34 (61.8%) | 7 (70%) |
| Female | 21 | 3 |
| R-ISS I | 15 | 1 |
| R-ISS II | 24 | 6 |
| R-ISS III | 4 | 2 |
| R-ISS unknown | 12 | 1 |
| M-component (median) (g/dl) | 28.1 (0-69.8)# | 10.45 (0-53.8)# |
| % PC from biopsy (median) | 45% (5-90%)* | 67.5 % (10-95%) |
| FISH t(11:14) | 5 | 1 |
| FISH t(14:11) | 1 | 0 |
| FISH t(4:14) | 9 | 3 |
| FISH del17p | 6 | 1 |
| FISH 1q21+ | 6 | 0 |
| FISH split IgH | 2 | 0 |
| FISH del13q | 1 | 0 |
| no translocations/deletions | 26 | 3 |
| LDH U/L (median) | 164 (71-441)§ | 162.5(117-235)§ |
| B2-mikroglobulin mg/L (median) | 3.65 (1.7-14)¤ | 5.9 (2.8-24.9)¤ |
| (short) PFS<2 years | 25 | 4 |
| (long) PFS>2 years | 29 | 6 |

### 2 BD patients unknown, 2 noBD patients unknown.

*1 BD patient <10 % PCs, 1 BD patient unknown.

FISH data: 7 BD patients unknown and 2 noBD patients unknown.

§ 9 BD patients unknown, 2 noBD patients unknown

¤ 11 BD patients unknown, 2 noBD patients unknown

Supplementary Table 4: Patient characteristics at diagnosis in untreated MM patients, divided into short (<2 years) and long progression free survival

| Characteristics | Short PFS (n=29) | Long PFS (n=34) |
|---|---|---|
| Age (median) y | 67 (45-79) | 64 (34-74) |
| Sex (male) | 21 (72.4%) | 20 (58.8%) |
| Female | 8 | 14 |
| R-ISS I | 4 | 12 |
| R-ISS II | 13 | 14 |
| R-ISS III | 4 | 2 |
| R-ISS unknown | 7 | 6 |
| Bone disease upon diagnosis | 25 (86.2%) | 28 (82.4%) |
| M-component (median) (g/dl) | 32 (0-67.1)# | 19.2 (0-69.8)# |
| % PC from biopsy (median) | 50% (5-90%) | 45% (8-95%)* |
| FISH t(11:14) | 3 | 2 |
| FISH t(14:11) | 0 | 1 |
| FISH t(4:14) | 5 | 7 |
| FISH del17p | 4 | 3 |
| FISH 1q21+ | 3 | 3 |
| FISH split IgH | 1 | 1 |
| FISH del13q | 0 | 1 |
| no translocations/deletions | 14 | 15 |
| LDH U/L (median) | 175.5 (71-319)§ | 162 (84-441)§ |
| B2-mikroglobulin mg/L (median) | 4.4 (1.8-11.6)¤ | 3.3 (1.7-24.9)¤ |

### 2 short_PFS patients unknown, 2 long_PFS patients unknown.

FISH data: 3 short_PFS patients unknown, 7 long_PFS patients unknown.

* 1 long_PFS patient with <10% PCs

§ 4 short_PFS patients unknown, 5 long_PFS patients unknown.

¤ 6 short_PFS patients unknown, 5 long_PFS patients unknown.
